# Cortical visual prosthesis: a detailed large-scale simulation study

**DOI:** 10.1101/610378

**Authors:** Jan Antolik, Quentin Sabatier, Charlie Galle, Yves Frègnac, Ryad Benosman

## Abstract

Recent advances in applying optogenetics in primates initiated the development of light based prosthetic implants for sensory restoration. Thanks to being the most well explored cortical area that is readily accessible at the surface of the brain, vision restoration via direct optogenetic activation of primary visual cortex is one of the most promising early targets for a optogenetics based prosthetic program. However, two fundamental elements of the cortical optogenetic prosthesis remain unclear. First, the exact mechanisms of neural dynamics under direct cortical stimulation, especially in the context of living, active and functionally specific intra-cortical neural circuitry, is poorly understood. Second, we lack protocols for transformation of arbitrary visual stimuli into light activation patterns that would induce perception of the said stimulus by the subject. In this study we address these issues using a large-scale spiking neural network modeling strategy of high biological fidelity. We examine the relationship between specific spatial configuration of light delivered to cortex and the resulting spatio-temporal pattern of activity evoked in the simulated cortical circuitry. Using such virtual experiments, we design a protocol for translation of a specific set of stimuli to activation pattern of a matrix of light emitting elements and provide a detailed assessment of the resulting cortical activations with respect to the natural vision control condition. In this study we restrict our focus to the grating stimulus class, which are an ideal starting point for exploration due to their thoroughly characterized representation in V1 and well-defined information content. However, we also provide an outline of a straight-forward road-map for transforming this grating centric stimulation protocol towards general strategy capable of transforming arbitrary spatio-temporal visual stimulus to a spatio-temporal pattern of light, thus enabling vision restoration via optogenetic V1 activation.

## 1 Introduction

Major effort is being undertaken to develop prosthetic implants [48, 67] for alleviating blindness in the millions of people suffering this condition around the globe [48]. Devices targeting retina have so far come closest to accomplishing reliable restoration of visual function, with several implants commercially available or in clinical trials [27]. However, for number of conditions which render retina or optic tract not viable, targeting the next two stages of visual processing – lateral geniculate nuclues (LGN) and primary visual cortex (V1) – are the two next best options. While being attractive target for vision restoration due to its straightforward coding of visual scene, the small size and poor accessibility deep in the brain make LGN a challenging option for visual prosthesis [61]. On the other hand, the accessibility at the surface of the brain combined with favorable magnification factor and well understood coding [23] predispose V1 as the primary target for extra-retinal vision restoration.

The recent advances in applying optogenetics in primates raise the possibility of using light as stimulation medium in neuro-prosthetic implants. Under such strategy, a region of cortex is transfected with opsin from the Channelrhodopsin family, rendering the transfected cells excitable by light [30]. A matrix of light emitting elements (MLEE) is placed on the surface of the treated cortical region (see figure 1). Pattern of light induced by the MLEE elicits analogous pattern of activity in the cortex under the implant. The central hypothesis is that, providing the induced cortical activity is sufficiently close to the encoding of the given visual stimulus under normal vision, perception of the said stimulus is elicited. Due to the absence of mechanical damage to the cortex and active electric currents, and the availability of high-density MLEEs (*<* 10*µm* pitch), such optogenetic based strategy elevates the disadvantages of implants relying on direct electrical stimulation [32, 33, 54, 66], while giving hope for restoration of high resolution vision.

**Figure 1:**
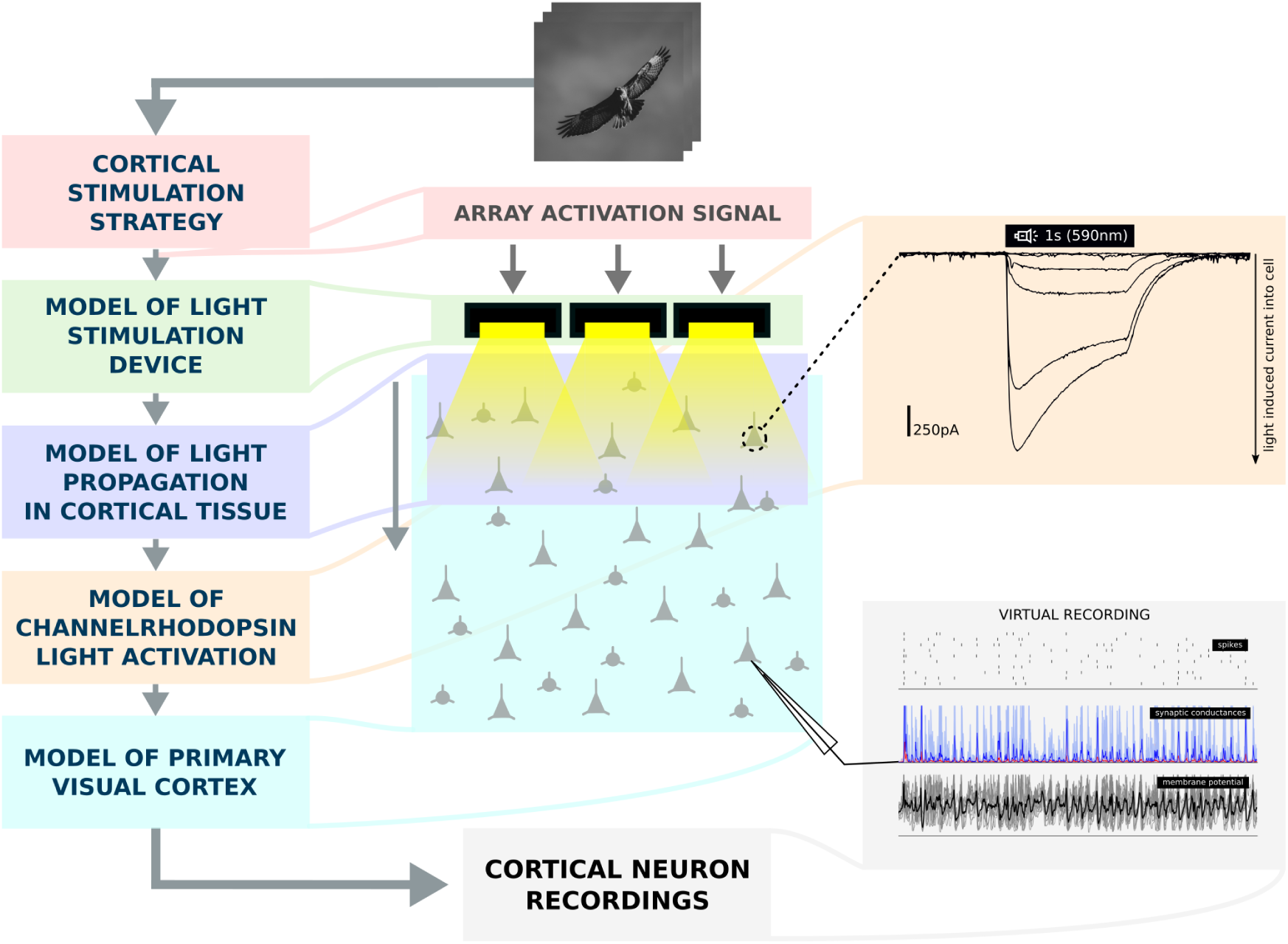
The schematic of the virtual cortical prosthesis experiment. (A) the stimulation strategy that translates a class of visual stimuli into driving signals for a MLEE, (B) a model of the MLEE (C) a model of light propagation through cortical tissue taking into consideration the absorption and diffraction of the light as it travels through the neural substrate. (D) a model of channelrhodopsin (ChR) dynamics in transfected cells that transforms a temporal trace of light impinging onto a given cell into a current that is injected into the cell due to the activation of ChR channels, and (E) a detailed large-scale spiking conductance based neural model of 5*mm*^2^ of primary visual cortex. Using these 5 simulation elements, we will be able to simulate the activation of cortical population to a specific set of visual stimuli. This in turn allows us to evaluate specific stimulation strategies by comparing the cortical activation patterns elicited by them to activation patterns due to equivalent stimulation via intact vision (through eye).

Two fundamental issues of such optogenetic visual prosthesis have, however, gained little attention so far. First, the mechanisms of cortical neural dynamics under external stimulation, especially in the context of living, active and functionally specific neural circuitry, is poorly understood. Second, we lack strategies for transforming arbitrary stimuli into light activation signals that would induce cortical activity patterns similar to those due to natural visual stimulation. Development of such stimulation strategies is challenging due to the complexity visual stimulus encoding schemes in V1 and the lack of deeper understanding of the interactions of light induced depolarization with the ongoing recurrent cortical dynamics. Therefore, computational models that capture both the complexity of the recurrent cortical dynamics and the V1 encoding of visual stimuli represent an ideal test bed for understanding the impact of light-activation on the cortical dynamics and in turn for design of light-based stimulation strategies.

Interestingly, surprisingly few computational studies have so far attempted to elucidate the impacts of optogenetic stimulation on cortical dynamics, and we are not aware of any studies trying to design an optimal light based stimulation protocol for eliciting activity patterns replicating the encoding of visual stimuli in V1. In this study we address both these issues using a detailed large-scale spiking neuron based modeling strategy of high biological fidelity. Specifically, using a combination of a model of light propagation in cortical tissue, model of channelrhodopsin dynamics, and a previous model of anatomically and functionally calibrated V1 cortical circuitry [10], we examine the cortical dynamics in a virtual population of transfected V1 neurons as a function of light stimulation parameters (figure 1). We demonstrate that performing these tests in a detailed recurrent model of cortex is crucial, as the simulated neural responses follow radically different patterns when the cortical circuitry is disabled.

Having gained insight on the dynamics of cortical activity under light stimulation, we proceed to formulate a light stimulation strategy for reproducing encoding of sinusoidal grating stimuli in V1. Focusing this canonical stimulus class with well studied encoding in V1 allows us to perform a systematic characterization of the light evoked neural dynamics and their similarity to analogous activity patterns evoked by natural visual stimulation via retina. We show that light activation of pyramidal cells in layer 2/3 at resolution of greater than 100 *µm* (in cortical coordinates), and at light intensities below 10^16^ *photons/s/cm*^2^, is sufficient to evoke cortical activity patterns close to those evoked by natural stimulation via retina. While here we only examine the canonical grating stimuli, in the discussion we provide a straightforward roadmap towards expanding this stimulation strategy to arbitrary spatio-temporal visual inputs.

To our best knowledge, this study represents the first quantitative examination of light based stimulation in functional model of cortical circuitry, and offers numerous predictions that can guide future experiments. Furthermore, we present the first V1 light stimulation protocol design that takes into consideration cortical dynamics under external stimulation, and thus provides a ready to use testbed for the upcoming in-vivo experiments of optogenetic prosthetic systems. Finally, the presented simulation framework is to our best knowledge the only currently available tool for simulation of the full cortico-prosthetic system, and can thus stimulate development int the sensory prosthetic field.

## 2 Materials and Methods

This manuscript relies on five key simulation components: (1) a stimulation strategy that translates a class of visual stimuli into driving signals for a MLEE, (2) a model of the MLEE, (3) a model of light propagation through cortical tissue, (4) a model of light illumination dependent channelrhodopsin (ChR) dynamics in transfected cells, and (5) a detailed large-scale model of primary visual cortex (whose parametrization and justification were developed elsewhere [10]). As illustrated in figure 1, the chaining of these five components allows us to simulate the effects of light stimulation in a ChR transfected primary visual cortex given a specific stimulation strategy. I.e. the visual stimulus is translated by the stimulation strategy into a set of signals determining the level of activation of individual light emitting elements. The light propagation model then determines the exact amount of light impinging onto individual neurons located in the simulated volume of cortical tissue given the pattern of activation of the MLEE. This in turn allows us to simulate the ChR dynamics for each cortical neuron given the amount of light it receives and thus determine the amount of current that it is injected with due to the light stimulation. Which finally allows us to simulate the behavior of the considered population of V1 neurons when embedded in the detailed, functional specific circuitry of the primary visual cortex.

This work has been based on our recent detailed large-scale model of primary visual cortex [10]. The model has been implemented using the Mozaik neural simulation workflow framework [8] and the Arkheia tool [7]. Here, we have extended the Mozaik framework with three additional components: the model of MLEE, the model of light propagation in cortical tissue, and a model of channelrhodopsin dynamics [65]. The NEST simulator [36] was used as the back-end for all simulations described in this paper. In the reminder of this section we provide a detailed description of the new components, and a more succinct description of the cortical model, referring reader to the previous literature for details.

### 2.1 The model of light emitting elements and light propagation in cortical tissue

We assume a regular 3.5 × 3.5*mm* lattice of circularly shaped light emitting elements of identical radius that is placed along the cortical surface. In this study we assume the pitch of the lattice to be 10*µm* and the individual elements having circular shape. This configuration corresponds well with for example a LED or DMD matrix commonly used in optogenetic stimulation. To determine the amount of light impinging onto individual neurons in the cortical volume situated under the matrix we performed two steps of simulation.

First we have determined the propagation of light through cortical tissue from a single light emitting element. We have performed the simulations using the ‘Human Brain Grey Matter’ model implemented in the LightTools software, assuming 590 *nm* wavelength of the emitted light. The scattering and absorption properties of the human brain tissue are modeled using the Henyey-Greenstein model [41]. The two key parameters of this model are the anisotropy factor *g* and mean free path (MFP) which are both dependent on wavelength. Considering the 590 *nm* wavelength we have set the two parameters to 0.87 and 0.07 mm based on Jacques *et al.*[41]. It should be noted that the accuracy and current knowledge of biological optical properties of cortical matter is limited and both the inter and within sample variability has been reported to be as much as 30%.

These simulations supply us with a 2D table *T* (*d, l*) capturing the light flux (*photons/s/cm*^2^) in cortical tissue relative to the value at the surface of the light emitting element as a function of depth *d* and the lateral distance (along the cortical surface) from the light source *l*. The light flux in tissue at a given level of activation of the light emitting element can be calculated as a simple multiplication of *T* by the light flux at the surface of the element. The light flux at the location of the given neuron *n* in the cortical volume is then calculated as a linear sum of the contributions from the individual elements in the matrix:

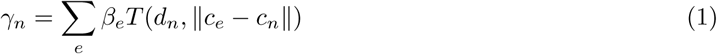

where *γ*_*n*_ is the resulting light flux at neuron *n, β*_*e*_ is the light flux at the surface of the light element *e*, and *c*_*e*_ and *c*_*n*_ are the lateral coordinates along the cortical surface of element *e* and neuron *n* respectively.

### 2.2 The channelrhodopsin model

We have used the model of ChrimsonR channel dynamics recently implemented by Sabatier *et al.*[65]. The electro-chemical behavior of the ChrimsonR protein is modeled using a Markov kinetic model [31]. In this model, a number of states (five in our case) represent the different conformations that the protein can take. For each pair of states, there can be a directed transition from one state to the other if there exist a chemical switch from the first state to the second. A time constant is associated with each transition. A transition can either be thermal or photo-induced. Thermal transitions have fixed time constants, while photo-induced transition’s time constants vary with the current value of the light stimulus. A photo-induced reaction cannot occur in the absence of light.

Mathematically, the values of the transitions time constants along with the light stimulus describe the linear differential system governing the evolution of the proportion of channels (or equivalently the probability for a single channel) in each state. The relevant figure, the conductance of the population of channels in a single neuron, is then derived from the number of channels in the open states and the conductances of these states.

The parameters of this model have been fitted by Sabatier *et al.*[65] to light (590nm wavelength) stimulation experiments in ChrimsonR-expressing HEK293 cells, and here we use the parameter values reported in that study.

### 2.3 The model of primary visual cortex

This model is derived from model presented in Antolik *et al.*[10]. A brief summary of the model follows. We refer the reader to Antolik *et al.*[10] for full details. The cortical model corresponds to layers 4 and 2/3 of a 3.5 *×* 3.5 mm patch of cat primary visual cortex, and thus given the magnification factor of 1 at 5 degrees of visual field eccentricity [73], covers roughly 3.5 *×* 3.5 degrees of visual field. It contains 24200 neurons and *∼*19 million synapses. This represents a significant down-sampling (∼10%) of the actual density of neurons present in the corresponding portion of cat cortex [14] and has been chosen to make the simulations computationally feasible. The neurons are distributed in equal quantities between the simulated layer 4 and layer 2/3, which is consistent with anatomical findings by Beaulieu & Colonnier [14] showing that in cat primary visual cortex approximately the same number of neurons inhabits these two cortical layers. Each simulated cortical layer contains one population of excitatory neurons (corresponding to spiny stellate neurons in Layer 4 and pyramidal neurons in Layer 2/3) and one population of inhibitory neurons (representing all subtypes of inhibitory interneurons) in the ratio 4:1 [15, 50].

We model both the feed-forward and recurrent V1 pathways; however, overall the model architecture is dominated by the intra-cortical connectivity, while thalamocortical synapses constitute less then 10% of the synaptic input to Layer 4 cells (see section 2.3.3), in line with experimental evidence [26]. The thalamic input reaches both excitatory and inhibitory neurons in Layer 4 (see Figure 2EF). In both cortical layers we implement short-range lateral connectivity between both excitatory and inhibitory neurons, and additionally in Layer 2/3 we also model long range excitatory connections onto other excitatory and inhibitory neurons [69, 22, 5] (see Figure 2AB). Layer 4 excitatory neurons send narrow projections to Layer 2/3 neurons (see Figure 2E). The model omits the infra-granular layer 5 and 6 as well as the cortical feedback to perigeniculate nucleus (PGN) and lateral geniculate nucleus (LGN).

**Figure 2:**
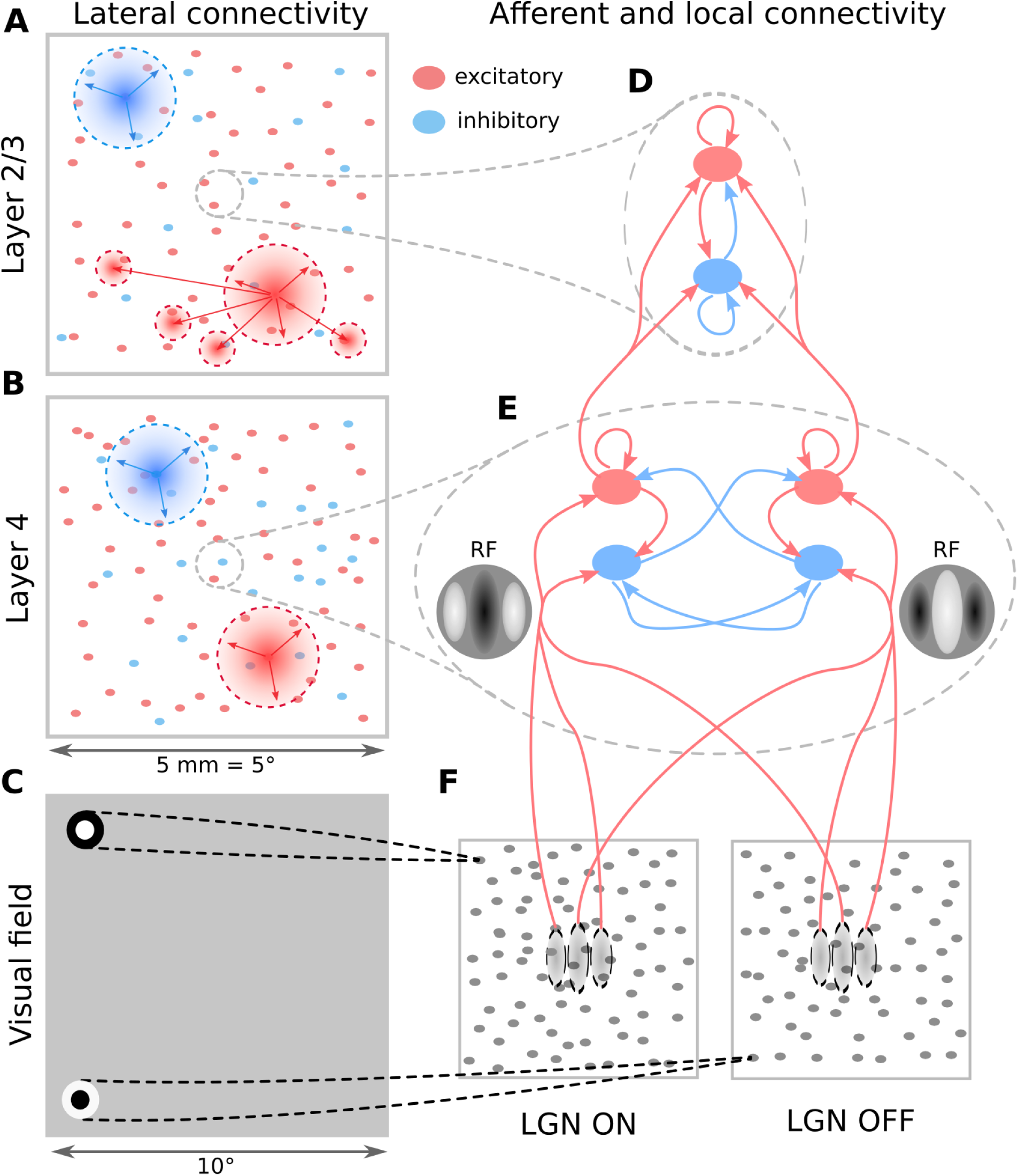
The model architecture. (A-B) Layer 2/3 and Layer 4 lateral connectivity. All cortical types make local connections within their layer. Layer 2/3 excitatory neurons also make long-range functionally specific connections. For the sake of clarity A,B do not show the functional specificity of local connections and connection ranges are not to scale. (C) Extent of modeled visual field and example of receptive fields (RFs) of one ON and one OFF-center LGN relay neuron. As indicated, the model is retinotopically organized. The extent of the modeled visual field is larger than the corresponding visuotopic area of modeled cortex in order to prevent clipping of LGN RFs. (D) Local connectivity scheme in Layer 2/3: connections are orientation-but not phase-specific, leading to predominantly Complex cell type RFs. Both neuron types receive narrow connections from Layer 4 excitatory neurons. (E) Local connectivity in Layer 4 follows a push-pull organization. (F) Afferent RFs of Layer 4 neurons are formed by sampling synapses from a probability distribution defined by a Gabor function overlaid on the ON and OFF LGN sheets, where positive parts of the Gabor function are overlaid on ON and negative on OFF-center sheets. The ON regions of RFs are shown in white, OFF regions in black.

#### 2.3.1 Neuron model

All neurons were modeled as the adaptive exponential integrate- and-fire units (eq 2), whereby the time course of the membrane potential *V* (*t*) is governed by:

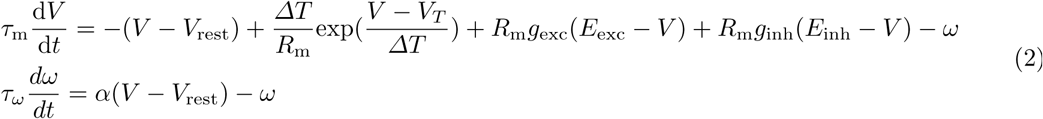

where *g*_exc_ and *g*_inh_ are the incoming excitatory and inhibitory synaptic conductances. Spikes are registered when the membrane potential crosses the 0 mV threshold, at which time the membrane potential is set to the reset value *V*_r_, and the spike-triggered adaptation mechanism is activated:

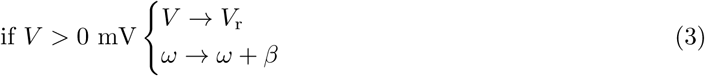

Each spike is followed by a refractory period during which the membrane potential is held at *V*_r_. All model parameters are set based on intra-cellular recordings of V1 neurons in Cat [53], and their values are listed in our previous manuscript [10].

#### 2.3.2 Thalamo-cortical model pathway

All neurons in the model Layer 4 receive connections from the model LGN (see Section 2.4). For each neuron, the spatial pattern of thalamo-cortical connectivity was determined by a Gabor distribution, inducing the elementary RF properties in Layer 4 neurons [72] (see Figure 2EF).

For individual neurons the orientation *θ*, phase *ψ*, size *σ*, frequency *λ* and aspect ratio *γ* of the Gabor distribution were selected as follows. To induce functional organization in the model, we used an existing model of stimulus dependent orientation map development [6] that utilizes Hebbian learning to compute stabilized link map that conditions an orientation map. Such pre-computed orientation map, corresponding to the 3.5 *×* 3.5 mm of simulated cortical area, was overlaid onto the modeled cortical surface, thereby assigning each neuron an orientation preference *θ*. The phase *ψ* of the Gabor distribution was asigned randomly, in line with the experimental evidence suggesting no clustering of spatial phase [77] in cat V1. For the sake of simplicity, the remaining parameters were set to constant values, matching the average of measurements in cat V1 RFs located in the para-foveal area [43], specifically the size *σ* was set to 0.5 degrees of visual field, the spatial frequency *λ* to 0.8 Hz and the aspect ratio *γ* to 1.7 [60].

#### 2.3.3 Cortico-cortical connectivity

The number of synaptic inputs per single excitatory neuron has been set to 800, which is about 25% of the actual number of synapses neurons in V1 receive on average [13]. This lower fraction has been chosen to exclude distal synapses that are not simulated in this model and due to reliability issues of synaptic transmission. For detailed justification please refer to Antolik *et al.*[10]. Inhibitory neurons received 35% fewer synapses than excitatory neurons to account for their smaller size, but otherwise synapses were formed proportionally to the two cell type densities. 30% of synapses from Layer 4 cells were formed on the Layer 2/3 neurons. In addition layer 4 cells received 80 additional thalamo-cortical synapses [26]. The synapses were drawn probabilistically with replacement (with functional and geometrical biases described bellow).

The geometry of the cortico-cortical connectivity was determined based on two main principles: the connection probability falls off with increasing cortical distance between neurons [20, 70, 22] (see Figure 2AB), and connections have a functionally specific bias, specifically they preferentially connect neurons with similar functional properties [22, 44]. The two principles were each expressed as a connection-probability density function, then multiplied and re-normalized to obtain the final connection probability profiles, from which the actual cortico-cortical synapses were drawn. The following two sections describe how the two probability density profiles of connectivity were obtained. Finally, apart from the connectivity directly derived from experimental data, we have also considered a direct feedback pathway from layer 2/3 to layer 4. Such direct connections from layer 4 to layer 2/3 are rare [16], however a strong feedback from layer 2/3 reaching layer 4 via layers 5 and 6 exists [16].

#### 2.3.4 Spatial extent of local intra-cortical connectivity

The exact parameters of the spatial extent of the model local connectivity, with the exception of excitatory lateral connections in Layer 2/3, were established based on a re-analysis of data from cat published in Stepanyants *et al.*[70]. For the exact description of this analysis and resulting parameter values please refer to Antolik *et al.*[10]. The Stepanyants *et al.*[70] study reflects only the local connectivity, due to it depending on neural reconstruction in slices which cut off distal dendrites and axons further than 500 *µ*m from the cell body. In cat Layer 2/3, unlike in Layer 4, excitatory neurons send long-range axons up to several millimetres away, synapsing onto other excitatory and inhibitory cells [5, 22]. To account for this long-range connectivity in Layer 2/3 we follow the observation by Buzás *et al.*[22], that the density of boutons across the cortical surface originating from the lateral connectivity of a small local population of stained Layer 2/3 neurons can be well approximated by the sum of two Gaussian distributions, one short range and isotropic and one long-range and orientation specific (see Section 2.3.5). Thus we model the lateral distribution of the Layer 2/3 excitatory connections as *G*(*σ*_*s*_) + *αG*(*σ*_*l*_), where *G* is a zero mean normal distribution, *σ*_*s*_ = 270 *µ*m and *σ*_*l*_ = 1000 *µ*m are the short and long-range space constants chosen in-line with Buzás *et al.*[22], and *α* = 1 is the ratio between the short-range and long-range components.

#### 2.3.5 Functionally specific connectivity

Within Layer 4 we assume push-pull connectivity [72] (see Figure 2E). For each pair of Layer 4 neurons the correlation *c* between their afferent RFs was calculated. The connectivity likelihood for a given pair of neurons is given by 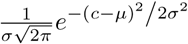 where *σ* = 0.3 and *µ* is 1 if the pre-synaptic neuron is excitatory or −1 if inhibitory.

In cat cortex excitatory neurons send long-range connections spanning up to 6 mm along the cortical distance to both excitatory and inhibitory neurons, preferentially targeting those with similar orientation preference [22]. To reflect this connectivity in the model we have defined the connectivity likelihood between pairs of neurons in Layer 2/3 as *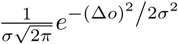* where the Δ*o* is the difference between the orientation preference of the two neurons, and *σ* was set to 0.3. Please refer to Antolik *et al.*[10] for detailed justification of the proposed connectivity scheme.

#### 2.3.6 Synapses

Synaptic inputs were modeled as transient conductance changes, with exponential decay with time-constant *τ*_*e*_ = 2.5 ms for excitatory synapses and *τ*_*i*_ = 6.0 ms for inhibitory synapses. We have set the unitary synaptic weight of excitatory synapses onto inhibitory neurons to 2.1 nS, while all remaining synapses in the model were set to 1.4 nS. We have also modeled synaptic depression for thalamo-cortical, and excitatory cortico-cortical synapses [1] using the model of Markram *et al.*[51], while we do not model short term plasticity for inhibitory synapses as it is not well studied. For the thalamo-cortical synapses we assume parameters corresponding to weak depression similar to Banitt *et al.* and Kremkow *et al.*[12, 46] (U=0.5,*τ*_rec_ = 120,*τ*_psc_ = 2.5 and *τ*_fac_ = 21). For the cortico-cortical excitatory synapses we have assumed stronger depression (U=0.5, *τ*_rec_ = 440 ms, *τ*_psc_ = 2.5 ms and *τ*_fac_ = 0 ms), in line with Markram *et al.*[51].

#### 2.3.7 Delays

We model two types of delays in the model circuitry. First are the delays due to the distance dependent propagation that are in the order of several tens of ms. These delays are important for lateral integration of information across multiple cortical columns. To reflect this, for all intra-cortical connectivity a distance-dependent delay with propagation constant of 0.3 ms^*-*1^ [19, 35, 42] was used, which corresponds to the slow propagation of action potentials along the intra-V1 (lateral) un-myelinated axons. The delays in the feed-forward thalamo-cortical pathway are drawn from a uniform distribution within the (1.4,2.4) ms^*-*1^ range. Second, Ohana *et al.*[56] have recently shown that delays of synaptic transmission in cat visual cortex are dependent on both pre- and post-synaptic neural type, with the notable feature of slow excitatory to excitatory and fast excitatory to inhibitory transmission. Distance-dependent axonal propagation delay is unlikely to explain these results as these experiments were performed in nearby neurons [56]. These pair-specific synaptic integration delays are in the order of only a few ms, but are important for local integration (within the same column) and for the precise timing of spike control by E/I interaction. Thus, as suggested by Ohana *et al.*, we have included a constant additive factor in all synaptic delays, specifically 1.4 ms for excitatory to excitatory synapses, 0.5 ms for excitatory to inhibitory synapses, 1.0 ms for inhibitory to excitatory synapses and 1.4 ms for inhibitory to inhibitory synapses, in line with the quantitative observations by Ohana *et al.*[56]. We observed that the addition of this neuron-type-dependent delay factor improved the stability of the modeled cortical neural networks, reducing synchronous events during spontaneous activity. We hypothesized that this is due to the ability of inhibition to respond faster to any transient increase in activity in the network due to the shorter excitatory to inhibitory delay.

### 2.4 Input model

The input model described below corresponds to the whole retino-thalamic pathway, where, for sake of simplification, retina and thalamus are treated as a single layer integration stage. Our cortical model corresponds to roughly 3.5 × 3.5*°* of visual field (Figure 2CF). To accommodate the full extents of RFs of neurons at the edges of the model, the LGN model corresponds to 4 × 4*°* of visual field. In the same manner, to accommodate the full extent of RFs of thalamic neurons the overall visual field from which the thalamic model receives input corresponds to 10 × 10°

We do not explicitly model the retinal circuitry and use the widely-used center-surround model of receptive fields (RFs) to simulate the responses of the LGN neurons (Figure 2C). The centers of both ON and OFF LGN neurons RFs are uniformly randomly distributed in the visual space, with density of 100 neurons per square degree. Each LGN neuron has a spatiotemporal receptive field, with a difference-of-Gaussians spatial profile and a bi-phasic temporal profile defined by a difference-of-Gamma-functions. Due to the relatively small region of visual space our model covers, we do not model the systematic changes in RF parameters with foveal eccentricity (nor, for the sake of simplicity, the natural cell-to-cell variance) and thus assume that all ON and OFF LGN neurons have identical parameters. The exact spatial and temporal parameters have been adopted from Allen and Freeman [4].

To obtain the spiking output of a given LGN neuron, the visual stimulus sampled into 7ms frames, was convolved with its spatiotemporal receptive field. In addition, saturation of the LGN responses with respect to local contrast and luminance is modeled [59, 18], please refer to Antolik *et al.* for details [10]. The resulting temporal traces are then summed and injected into integrate-and-fire neurons as a current, inducing stimulus dependent spiking responses. In addition to the stimulus-dependent drive, neurons are also injected with white noise current. The magnitude and variance of this noise is such that neurons fire *∼* 10 spikes/s in the no stimulus condition [72]. This artificially elicited spontaneous discharge, that is calibrated to reproduce the experimentally observed spontaneous rates, corresponds to the combined effects of the dark discharge of the retina and any other potential intrinsic mechanism of spontaneous activity generation in the thalamus.

### 2.5 The in-silico experimental protocols

Throughout this study we employ orientation tuning protocol, that uses the drifting sinusoidal grating stimulus of varying orientation to probe the canonical functional property of V1 neurons: the orientation preference and selectivity. We administer this protocol in two variants, first that simulates the stimulation of the cortex via retina (normal vision condition) and second that simulates the replication of the same stimulation protocol via light activation (the prosthetic vision condition). The orientation tuning protocol consists of series of sinusoidal grating stimuli that are presented at 8 different orientations in equal steps between 0 and 180 design. Each grating was shown 10times for 600 ms. The spatial and temporal frequency of the RFs of the modeled LGN neurons (see Section 2.3.2) and of the Gabor distribution template from which thalamo-cortical synapses were sampled were identical. An important consequence of this simplification is that it allowed us to efficiently execute protocols requiring drifting sinusoidal gratings. By employing a full-field stimulus with spatial frequency matching that of the Gabor template (0.8 Hz) and drifting at 2 Hz, we were in parallel stimulating all cortical neurons with a stimulus with optimal spatial and temporal frequency.

Section 3.1 describes light stimulation protocol for evocation of cortical activity corresponding to visual stimulation by specific full-field sinusoidal grating stimulus. Given this, the prosthetic vision variant of the orientation tuning protocol followed the same series of corresponding stimuli as described in previous paragraph, rendering the evoked responses directly comparable between the two tuning protocol variants.

In order to assess orientation tuning of the responses resulting from the tuning protocol described above (in both natural vision and prosthetic conditions), we followed [55] and calculated the half width at half height (HWHH) measured by fitting the orientation tuning curves with a Gaussian function [3]:

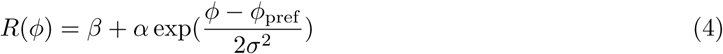

where *R* is the spiking response of the given neuron to a sinusoidal grating with orientation *ϕ, ϕ*_pref_ is the preferred orientation of the given neutron, *σ* is the width of the tuning, *β* is the baseline activity and *α* a scale factor. Low responding neurons (less then 1 spike/s at optimal orientation) were excluded from the analysis, as reliable curve fitting was not possible with the amount of recorded data. Furthermore, neurons (*<* 10%), for which reliable fit of Gaussian curve was not possible (*MSE >* 3 0 % of the tuning curve variance) were also excluded from this analysis. HWHH was then calculated as *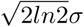*.

### 2.6 Data analysis

We have used the Naka-Rushton function to fit the contrast-response (figure 6ABC) and light intensity-response (figure 6E) relationships:

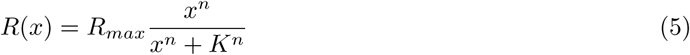

where *R*(*x*) is the response at either contrast level *x* or stimulation light intensity *x, R*_*max*_ is the asymptotic maximum response amplitude, *K* is the semi-saturation constant, and *n* controls the slope.

In Figure 5D we fit the relationship between the photon flux at the cell body and the response of the cell with a sigmoid:

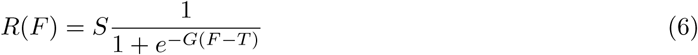

**Figure 3:**
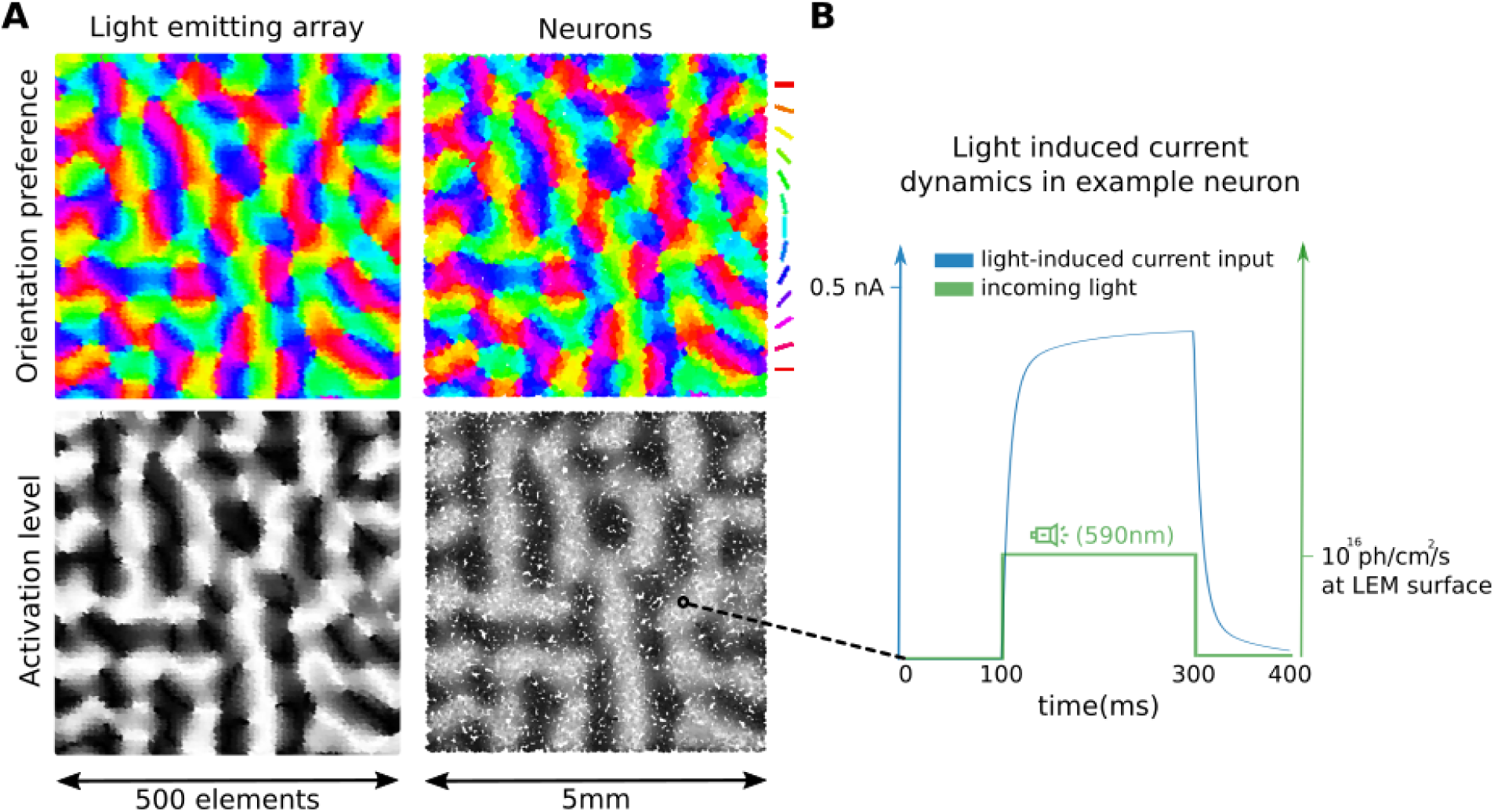
Opto-stimulation protocol for grating stimulus. (A) The orientation preference of neurons and assigned orientation preference to individual light emitting elements (top row); The driving signal to individual light-emitting elements determined by the proposed stimulation protocol and the resulting depolarization in cortical tissue (bottom row). (B) The light output of an example light-emitting element (green) and the resulting light-mediated inward current to an example neuron located at the same location (blue).

**Figure 4:**
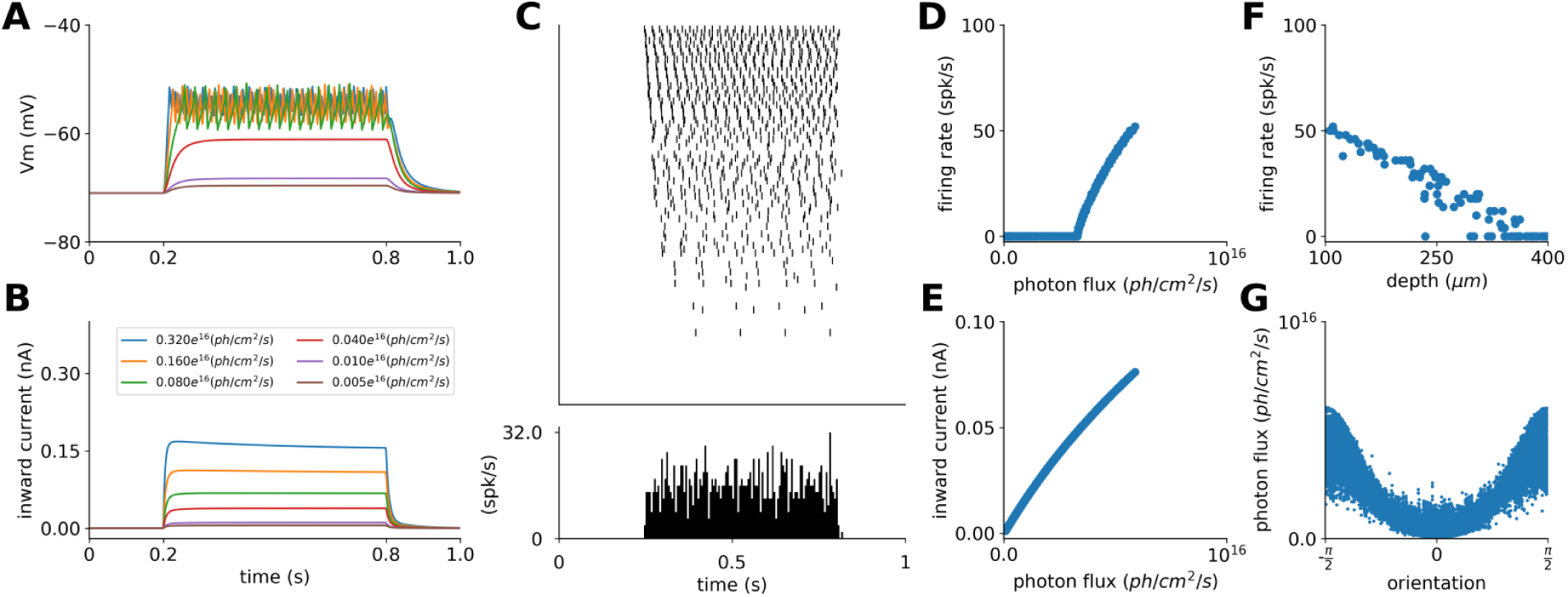
Light stimulation in cortical model. (A) The membrane potential of a randomly selected cell at different levels of intensity of light impinging on the cell (the color coding corresponds to panel B). The light stimulation was a step function, starting at 100ms and ending at 800ms.(B) The inward current elicited by the light stimulation of different intensity (the legend shows the maximum photon-flux at the surface of the MLEE) in the same cell as in A. (C) The spiking response of population of cells recorded in central region of the model in response to light stimulation. The cells were ordered according to their cortical depth increasing from top to bottom. (D) The relationship between the photon flux at the position of neurons in the cortex and their firing rate. (E) The relationship between the photon flux at the position of neurons in the cortex and the resulting inward current. (F) The relationship between the depth of neurons in the model cortical substrate, and their response. (G) The relationship between orientation preference of neurons (abscissa) and their firing-rate response (ordinate) to light stimulation emulating sinusoidal grating of 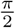 orientation. The stimulation followed the strategy presented in section 3.1. With the exception of (G) only neurons whose orientation preference matched with the orientation of the light-stimulation emulated grating stimulus were considered.

**Figure 5:**
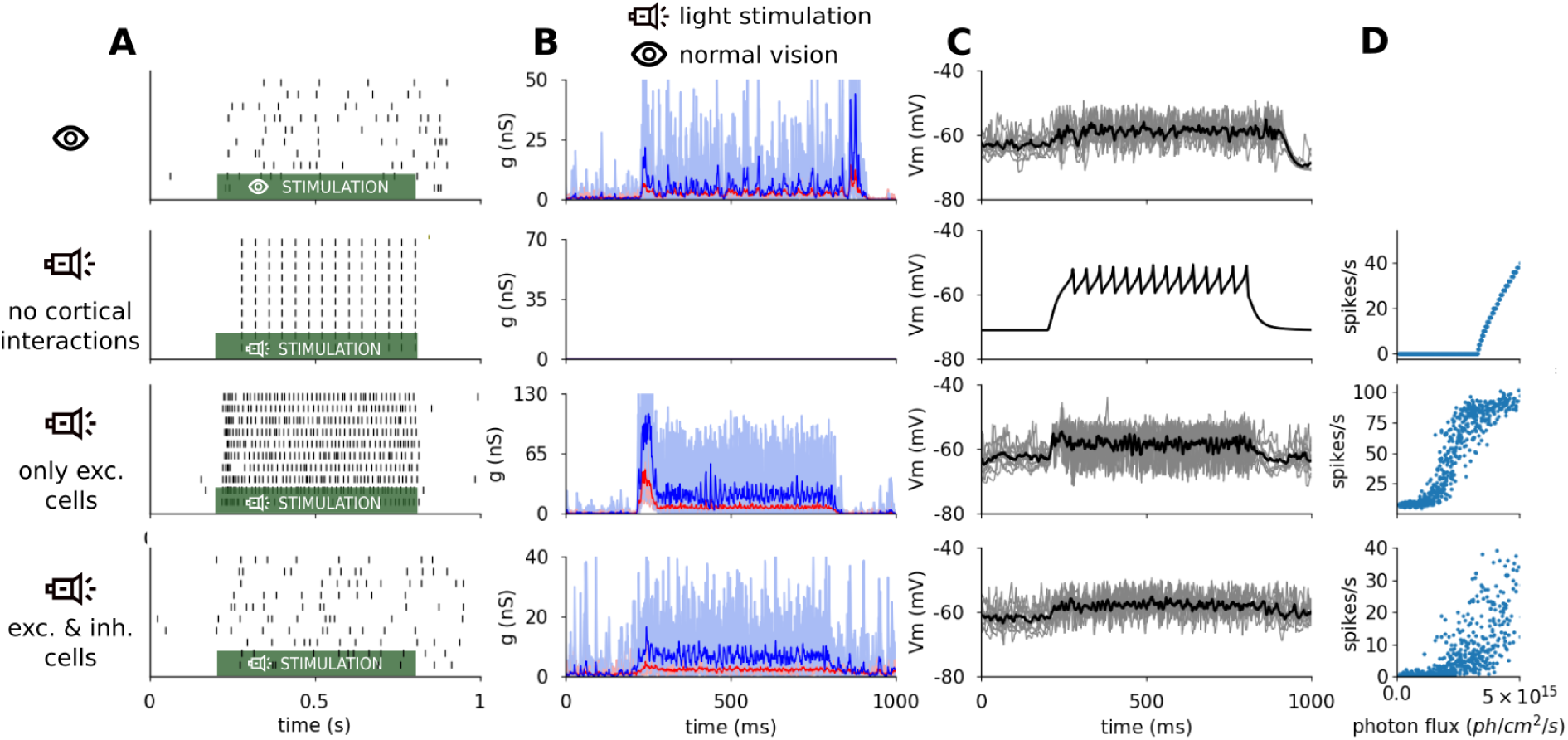
Example single-neuron dynamics in the 4 experimental conditions. The neurons respond to 10 trials of stimulation with drifting sinusoidal grating (or its emulation via cortical proshesis), that starts at 100ms and stops at 800ms. From top to bottom, intact system with stimulation via retina, cortical light stimulation with all model connectivity disabled, cortical light stimulation where only excitatory neurons respond to light, and cortical light stimulation where both excitatory and inhibitory neurons respond to light. (A) spike raster plot. (B) Excitatory (red) and inhibitory (blue) conductances. (C) Membrane potential. (D) The relationship between photon-flux and the spiking response of neurons in the given condition. In B and C light thin lines are single-trials, thick saturated lines are mean across trials. In all cortical stimulation conditions, the stimulation scaler was *L*_*max*_ = 9.2 × 10^15^ *photons/s/cm*^2^.

*R*(*F*) is the response at photon flux *F*, *S* is the scaler, *G* is the gain, and *T* is the threshold parameter. The threshold *T* parameter of the sigmoid function doesn’t reflect well the threshold of the stimulus-response relationship that is typically derived in experimental setting as the point where the response function departs significantly from the spontaneous rate. To obtain a more comparable quantity we have calculated what we will refer to as effective threshold (eT), which is the photon-flux where the fitting sigmoid reaches 0.05% of its asymptotic maximum.

## 3 Results

The central goal of this study is to use computational methods to compare the dynamics of population of layer 2/3 neurons within a local volume of primary visual cortex under natural visual stimulation condition against the condition where the cortical representation of the same stimulus are approximated via opto-genetically mediated light stimulation using matrix of light emitting elements. In this study we will restrict ourselves to the experimentally most extensively characterized stimulus class: the drifting grating stimuli. We will perform a series of virtual experiments in which we will explore several parameters of the grating stimulation protocol, gradually improving the match between visually and light induced cortical responses.

### 3.1 The stimulation strategy

Because the drifting grating family of stimuli are effective drivers of activity in primary visual cortex, while being mathematically highly tractable, they became the dominant visual stimulation method for studying V1. Consequently, the behavior of V1 neurons with respect to the changes of the parameters of the grating stimulus (such as orientation, contrast, frequency, etc.) became probably the most well characterized aspect of cortical visual processing [39, 64, 29]. Here we will restrict our attention to this stimulus class, designing a light stimulation strategy that can elicit cortical responses reminiscent of those evoked by drifting gratings. However, using insights from the presented work, in the Discussion we offer an outline of a straightforward method to extended this stimulation strategy to arbitrary spatio-temporal stimulus.

Majority of neurons in V1 are selective to the orientation of the stimulus [24]. With respect to phase of the grating, two functional cell classes have been identified [37]. Neurons in the first class respond linearly whereby only a specific phase of the grating elicits their spiking response - the so called simple cells. Neurons in the second class respond non-linearly, firing spikes to any phase of the grating, and thus they respond by a tonic elevation of their membrane potential and spike response to a drifting grating that is superimposed over their receptive field - the so called complex cells. These two functional cell types are not distributed randomly throughout the cortical layers, but simple cells occupy predominantly layer 4 while complex cells form the majority of cells in layer 2/3 [38, 64]. In the horizontal plane along cortical surface, the functional properties of neurons also follow specific organization, whereby nearby cells tend to be selective to similar orientation of the stimulus [40]. When viewed from the surface of the cortex, the orientation preference of cells thus forms the signature orientation maps with smoothly varying orientation preferences regularly interrupted by local discontinuities, so called pinwheels [25].

Due to the absorption and dispersion of the light as it travels through cortical tissue, the intensity and resolution (contrast) of the pattern of light induced by the array of light emitting sources degrades with increasing cortical depth (see section 2.1). Layer 2/3 being the primary layer that sends axons to higher cortical areas (i.e. the cortical output layer), while at the same time being close to its surface is thus the ideal target for optogenetic based vision restoration intervention.

Taking into consideration all the above anatomical, functional and physical constraints, we propose following stimulation strategy:

1. Assuming the knowledge of orientation preference map in the targeted cortical volume, assign orientation preference *OR*_*M*_ to each light emitting element *M* located at the cortical surface coordinates *C*_*M*_ based on the circular distance weighted average of orientation preferences of neurons centered at coordinates *C*_*M*_ (figure 3A).

2. For a full-field sinusoidal grating of orientation *ρ*, for each light emitting element *M* calculate the orientation dependent activation index *ψ*_*ρ,M*_ as *ψ*_*ρ,M*_ = *f* (*δ*(*ρ, C*_*M*_)), where *δ* is circular distance and *f* is a function of distance. In this study we will set *f* to a Gaussian function with zero mean and *σ* = 0.5 variance (except in section 3.3 where *σ* is varied; figure 3A).

3. Set the signal driving light emitting element *M* as a step function such that the resulting light output at its surface (photon flux measured in *photons/s/cm*^2^) is *θ*_*M*_ = *S*_*M*_ (*τ*_*s*_, *τ*_*e*_, *ϕ*), where *τ*_*s*_ is the start of the step, and should be set to the delay of layer 2/3 activation in V1 by grating stimulus under intact vision condition, and the *τ*_*e*_ = *τ*_*s*_ + *d* is the end of the step, where *d* is the duration of the grating stimulus (figure 3B). The *ϕ* = *L*_*max*_*ψ*_*ρ,M*_ is the magnitude of the step, where *L*_*max*_ is an overall stimulation scaling factor setting the maximum light emission at the surface of the MLEE (i.e. the *L*_*max*_ level of photon flux will be achieved only at the surface of the light emitting elements that perfectly match the orientation of the to be induced grating stimulus).

The *L*_*max*_ is an arbitrary scaling factor that has to be determined experimentally to achieve desired level of activation, and absorbs such scaling unknowns as the rate of ChR transfection of the given cortical volume, or the intensity of light emission of the matrix element depending on the driving signal. Importantly, it also takes into account stimulus dependent scaling. In the case of sinusoidal gratings, it is the contrast of the stimulus that sets the overall magnitude of membrane potential depolarization, and hence the tonic level of spiking response. In section 3.3 we will offer a simple methods how to determine the *L*_*max*_ parameter in a stimulus contrast dependent manner. Finally note that in principle the *L*_*max*_ also depends on other properties of the grating stimulus to which V1 neurons are selective, such as its spatial or temporal frequency, but for the sake of simplicity in this study we will assume that the other grating parameters are kept at the known preferred values at the given retinotopic eccentricity of the targeted V1 volume, and the *L*_*max*_ is being determined with respect to these fixed values.

### 3.2 In-silico opto-stimulation of primary visual cortex

To demonstrate the integrity of the simulation platform and to gain basic intuition on the elementary influences of light stimulation on the cortical substrate, we will first examine a version of the model in which all connections between the neurons were disabled. In other words, all cortical model neurons behave independently of the other neurons, and, because we do not assume any external source of noise in the model, their membrane potential is fully determined by only two factors: (i) their position in the cortical volume and (ii) the induced pattern of light.

We have stimulated such ‘disconnected’ model with a light stimulation pattern designed to approximate visual stimulation with drifting sinusoidal grating of horizontal orientation and optimal spatial and temporal frequency, as described in previous section. In figure 4 we show responses of model neurons to such stimulation. With the exception of panel G, only neurons with preference for horizontal orientation, matching the orientation of stimulating grating, are analyzed. Figure 4A shows the evolution of membrane potential of an example layer 2/3 model cell in response to step light stimulation pulse of different intensity. The inward current to the same cell due to the same light intensities are shown in figure 4B (color-matched traces between A and B). The traces in Fig4B bare the characteristics of the ChR dynamics (see Methods), which are further transformed by the membrane potential dynamics as shown in figure 4A. As expected, increased intensity of light stimulation leads to increased inward current due to activation of ChR (Fig4E), which in turn leads to increasingly depolarized membrane-potential and once the action potential threshold is crossed, light intensity is correlated with firing rate (figure 4D). Another way to view this is in figure 4C, which shows spike raster plot for a population of randomly selected model neurons, one line per neuron, which were ordered by the depth of their location in the cortex. We can see, that as the cortical depth of the neuron increases, their response rate decreases, as expected due to the attenuation of incoming light which is the result of absorption and dispersion of light as it travels through the neural substrate. This depth-dependence of firing rate is more clearly visualized in figure 4F. In figure 4C we can also notice a slight increase in onset time of the response with increasing depth, which is due to the reduced driving force causing slower and thus longer evolution from resting to threshold membrane potential that induces the first action potential. Finally, figure 4G shows the photon flux as a function of the orientation of the neuron, clearly demonstrating the desired orientation dependence of stimulation. We will examine this orientation tuning more closely in section 4. Finally, we would like to note, that due to the absence of any ongoing noise in the decoupled model all the variations in figure 4F and G are due to the variations of neuron’s depth or lateral displacement of neurons with respect to the orientation map features (see Methods).

Having verified the sanity of the basic simulation properties in the decoupled model, we can proceed to explore the response properties of cells under the dynamics induced by the fully connected cortical circuit. In the remainder of this study we will consider 4 basic conditions comparison of which will facilitate understanding of the simulation outcomes: (i) the model stimulated with the gratings stimulus via retino-thalamic pathway (i.e. the natural vision condition), (ii) an opoto-genetically stimulated decoupled model, (iii) an opoto-genetically stimulated fully connected model where only excitatory cells express Channelrhodhopsin and (iv) an opoto-genetically stimulated fully connected model where both excitatory and inhibitory neurons express Channelrhodhopsin.

Figure 5 shows the response of an example neuron to 600ms presentation of horizontal sinusoidal grating or its opto-genetically induced equivalent. In the natural vision condition the grating is presented at 100% contrast, while for all the opto-genetically stimulated conditions the same arbitrary magnitude of stimulation (*L*_*max*_ = 9.2 × 10^15^ *photons/s/cm*^2^) was selected such that all conditions induce sufficiently elevated firing response (the problem of matching the scale of opto-genetic activation with stimulus contrast will be addressed in the next section). As shown in figure 5, neurons in all conditions respond to the stimulation by a tonic elevation of the membrane potential and spiking response for the duration of the stimulus, but beyond these basic characteristic number of clear differences are present. The membrane potential and conductances in the natural vision condition exhibit clear transient on- and off-set dynamics due to the thalamic and cortical processing (see Methods), in line with experimental evidence [66, 49, 53]. Since the proposed stimulation protocol does not explicitly address these temporal response characteristics and because under normal vision they arise largely due to feed-forward mechanisms [52] which aren’t engaged during cortical stimulation condition, it is not surprising that the optogenetical stimulation conditions lack them. Next, one can observe difference in the overall level of membrane potential depolarization, its variance as well as the absolute magnitude and ratio of the excitatory and inhibitory conductances, which will be examined in detail in section 5.

Last but not least, we can notice major differences in the overall firing response. Among the optogenetically stimulated conditions the reasons for these differences are straightforward: in the decoupled condition no inhibition and source of noise are present and the condition thus exhibits steep illumination-spiking curve (figure 5D), that is offset to the right due to the difference between resting membrane potential and spiking threshold that has to be crossed by light induced depolarization before any spikes are fired (G=2.62, eT=3.00 × 10^15^ *photons/s/cm*^2^, see Methods 2.6). Even though inhibition is present in the coupled condition with only excitatory neurons activated by light, steep activation curve reaching higher values is observed and the intensity threshold required to induce firing becomes lower (G=1.82, eT=0.51 × 10^15^ *photons/s/cm*^2^, see Methods 2.6). This difference can be explained by the stochasticity of the membrane potential due to the ongoing activity in the coupled model. Due to the variability of the membrane potential even small extra light-induced depolarization will generate extra spikes, shifting the illumination-spiking curve to the left, and thus smoothing the transition between no-response and activated intervals. Furthermore, the stochasticity of the membrane potential means that the same levels of mean depolarization as in the decoupled condition will result in higher response rates in the coupled condition. Finally, in the third opto-genetic condition with both excitatory and inhibitory neurons activated by light, we can observe lower gain and higher threshold of the illumination-spiking curve (G=1.02, eT=1.71 × 10^15^ *photons/s/cm*^2^, see Methods 2.6), due to the additional external inhibitory drive shifting the E/I ratio in favor of the inhibition.

These differences in overall response magnitudes across conditions pose two problems: (i) for the purpose of effective prosthetic stimulation protocol we need to be able to control the magnitude of response and match it to physiological levels expected for the given visual stimulus, and (ii) for the purpose of this study it challenges our ability of doing rigorous comparisons across conditions as many neural signal measures are rate dependent. We will address these issues in the following section, but before we continue, let us point out that these basic observations of single-cell model responses already have important implications for cortical optogenetic stimulation. First, they demonstrate that considering the neural network dynamics resulting from the intra-cortical connectivity is essential for drawing relevant conclusion that can drive future experiment. Second, extrapolating optogenetic single cell results whether from modeling, cell culture or slice experiments that all largely lack ongoing activity and network effects can be problematic. In particular, these observations suggest, that optogenetic experiments in slices or cultures will over-estimate the minimum amount of light necessary to evoke spike-response in in-vivo conditions (as long as other experimental parameters are well matched between the in-vivo and in-vitro conditions). In our hands, this overestimation is about 6-fold (eT=3.0 (uncoupled condition) vs eT=0.51 (coupled condition selective to excitatory cells)). We also predict that the excitatory to inhibitory cell type specificity of ChR transfection will have major impact on the intensity of light required to induce desired neural response levels, and will effect the operating regime of the network.

### 3.3 Calibration of contrast response curves

In complex cells of primary visual cortex, stimulation with drifting sinusoidal grating induces steady depolarization of membrane potential leading to a tonic increase in spiking of the cell. The magnitude of the depolarization and in turn the neuron’s firing rate grows with the contrast of the stimulus. Because under the light stimulation the membrane potential depolarization is proportional to the intensity of the incoming light, in the proposed stimulation paradigm the analogue of the contrast parameter is the light intensity scaler parameter *L*_*max*_ (Methods 3.1). However, both the relationship between the contrast and response rate, and the light intensity and response rate is non-linear (see figure 4). In what follows we quantify these input intensity-response relationships in the model, and demonstrate a scheme for mapping contrast of a grating stimulus onto light intensity that induces matching magnitude of response. This allows us in the reminder of the manuscript to perform comparisons between visual and light based stimulation that are matched for the overall response magnitude.

Figure 6A-C shows light intensity to response curves of example model layer 2/3 cells and corresponding averages across the measured population for the 3 examined optogenetic conditions. As expected, in the visual stimulation condition (figure 6E black line) the contrast-response curve is well fitted with the so called Naka-Rushton function (see methods) as demonstrated in number of in-vivo studies [34]. Conveniently, the Naka-Rushton function also fits well the relationship between the light intensity and evoked firing rates in the 3 optogenetic stimulation conditions (figure 6A,B,C). Just as in-vivo neurons, our model neurons exhibit considerable variability in the fitted parameters of the Naka-Rushton function from cell to cell. Furthermore, in the optogenetic conditions, there are systematic changes with respect to depth due to the light absorption and dispersion in cortical substrate. Due to the various constraints of the optical cortical prosthesis approach assumed here, it is impossible to control the activity of each individual neuron independently, rather in our stimulation protocol we have only one global parameter - the light intensity scaler *L*_*max*_ - to control the global level of depolarization of neurons in the targeted cortical volume. Therefore, the best approximation we can hope to achieve is to match the mean contrast-response curve across the targeted cortical volume. Having the parametric fits of the light-to-response tuning curve *F* and contrast-to-response tuning curve *G*, this can be achieved by simply mapping the desired contrast onto rate using *G* and then mapping this rate using inversion of *F* on the desired light intensity scaler. Thus the function composition *F* ^*-*1^(*G*) maps the contrast of the stimulus to be induced onto the light intensity scaler that will induce it (figure 6C). As is demonstrated in figure (figure 6D), this method secures a good match of the mean contrast response curves between the natural vision and optogenetic conditions. In the remainder of this study, we will be comparing the different stimulation conditions through the contrast matched paradigm demonstrated in this section.

## 4 Light induced orientation tuning of cortical responses

We will now turn our attention to the most salient and well explored property of V1 neurons: their orientation selectivity. Majority of neurons in V1 are selective to orientation of the stimulus, such that they response is highest when stimulus orientation matches their orientation preference, and response drops of as the orientation of the stimulus departures from the preferred orientation [39]. V1 neurons rarely respond with any spikes (beyond spontaneous level) to stimuli orthogonal to their preferred orientation [34]. The tuning curves formed by such orientation specific responses can be well fit with Gaussian curves, and selectivity in majority of neurons is sharp, falling between 15 and 30 degrees when measured as the half-width at half height (HWHH) of the fitted Gaussian [55, 24, 34]. Another important property of the orientation selectivity in V1, that cannot be explained in a simple feed-forward model, is that the width of their tuning curve is independent of contrast - i.e. changes to contrast only induce multiplicative changes to the response.

The cortical circuit model upon which the present study is derived from has been extensively explored with respect to its functional characteristics and has been show to match all the above properties (and many others) [10]. Indeed when we repeat the orientation selectivity experiments in the natural vision condition in our simulation environment, we find that most neurons express Gaussian shaped (figure 7A) contrast invariant (figure 7B) orientation tuning curves with mean tuning width of about 25 degrees (see figure 7C).

**Figure 6:**
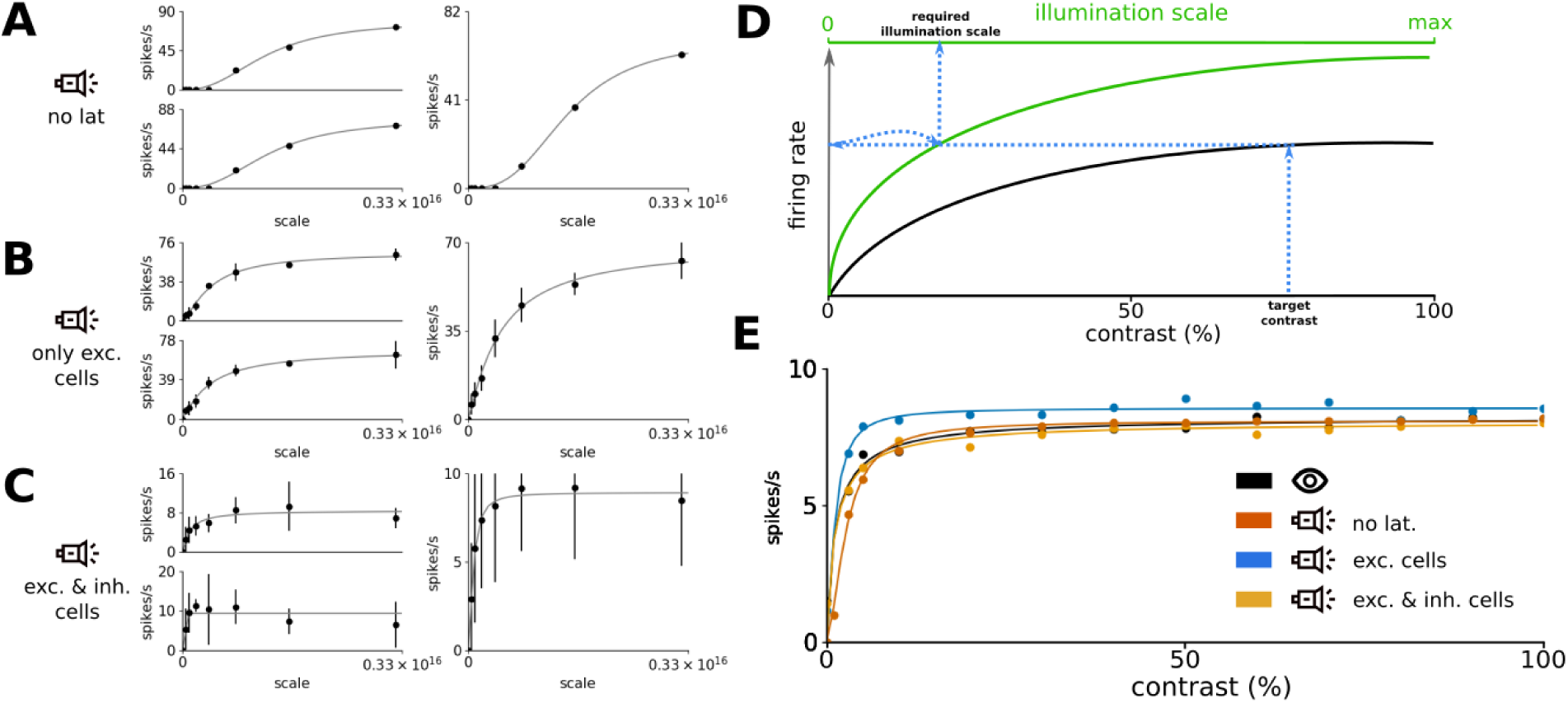
Contrast to stimulation light intensity mapping. (A-C) The stimulation light to response curve of two example individual neurons (left panel) and the mean curve over all recorded neurons (right panel). Three conditions shown: no lateral connectivity (A), only-excitatory neurons excitable by light (B) and both excitatory and inhibitory neurons excitable by light (C). (D) A scheme for translation of desired contrast of the to be induced grating stimulus to the corresponding required stimulation light intensity. (E) The resulting contrast-response curves of the three optogenetic conditions superimposed over the natural vision data. In all panels, the dots are data points from simulations, line is the fit of the Naka-Rushton curve to the data points.

**Figure 7:**
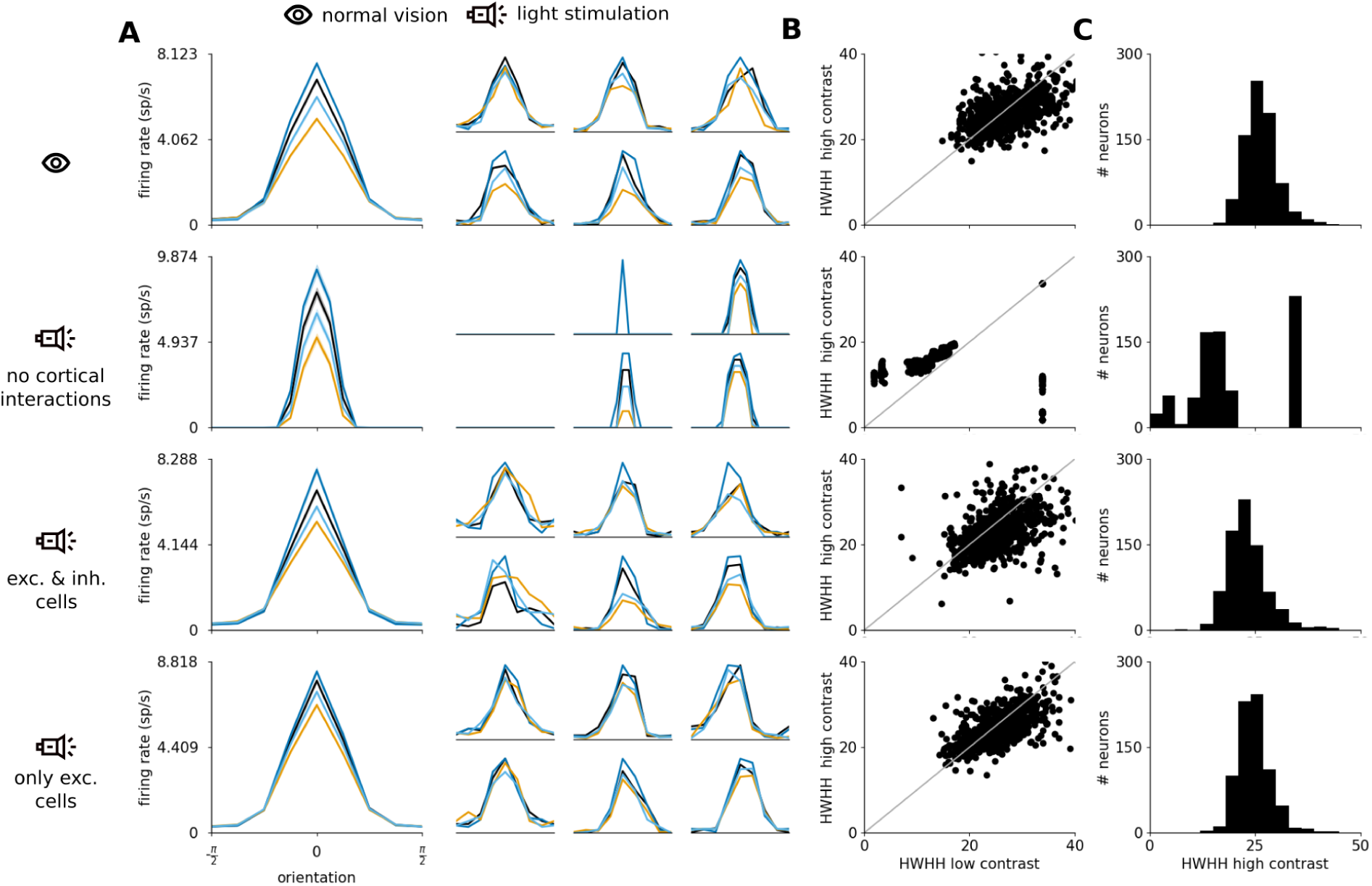
Orientation tuning of model neurons in the 4 experimental conditions. From top to bottom, intact system with stimulation via retina, cortical light stimulation with all model connectivity disabled, cortical light stimulation where both excitatory and inhibitory neurons respond to light, and cortical light stimulation where only excitatory neurons respond to light. (A) mean of centered orientation tuning curves across all recorded neurons (large plot) and single cell tuning curves of 6 randomly selected neurons (6 small plots). (B) the half-height at half width (HWHH) of orientation tuning curves of neurons when stimulated at low contrast or maximum light intensity (abscissa) and high contrast or maximum light intensity (ordinate). (C) the distribution of HWHH at high contrast/light intensity across the population of measured neurons.

Having functionally accurate benchmark in the natural vision condition, we have repeated the same orientation tuning protocol but using the analogous optogenetic stimulation (see section 3.1) in the decoupled model. In this condition the model response directly reflects the orientation selectivity of the stimulation protocol (i.e. see the parameter *σ* in section 3.1), only altered by the light propagation properties of cortical tissue, but not shaped by the intra-cortical connectivity. We can see that while the responses are clearly orientation tuned, their broadness increases with contrast of the stimulus (see figure 7A,B). Importantly, by increasing light intensity further would broaden the tuning curves, eventually eliciting responses even in neurons with orthogonal orientation preferences to that of the stimulus (figure S1). This is not the fault of the stimulation protocol itself: the light emitting elements sitting above the cortical domains with orthogonal orientation preference are set to be off by the stimulation protocol (see section 3.1). Instead it is two opposing phenomena that cause the contrast dependent changes in the width of the tuning. On one hand it is the dispersion of light traveling along the cortical tissue that causes bleeding of light from those domains that are being illuminated into nearby domains that are not meant to be illuminated. It is this phenomena that can cause, providing sufficiently intense illumination, even invocation of activity in neurons that prefer orthogonal orientation. On the other hand, due to the lack of ongoing activity in the decoupled model, the gap between the resting and membrane potential and spiking threshold causes an ‘iceberg effect’ whereby any cortical volume that is not illuminated above fixed threshold will generated zero response. It is this second phenomena that in the range of the illuminations we test here (dictated by the contrast response curve calibration discussed in section 3.3) causes the very narrow tuning curves that abruptly cross zero response level at about 45 degrees, unlike the gradual decrease towards zero of the natural vision condition.

Note that while for fixed contrast one could possibly tune the stimulation protocol such that the right tuning is achieved in the decoupled condition it could not be done in a contrast independent manner. Could however the engagement of intra-cortical connectivity that is know to exert competitive influence over the cortical response, whereby stronger responses are magnified, while weaker suppressed resolve these difficulties we observe in the decoupled condition? Indeed in both optogenetic conditions with intact intra-cortical interactions the model neurons exhibit orientation tuning curves remarkably similar to those due to visual stimulation (figure 7): both optogenetic conditions generate tuning curves with similar tuning width and contrast invariance. We would like to point out that this is an interesting and encouraging finding. While conceptually the presence of complex recurrent dynamics due to the recurrent connectivity complicates the reasoning how to optimally stimulate the cortex, these results suggest that their presence can actually make it easier (or even possible at all) to optogenetically induce cortical activity patterns that are close to those due to natural vision.

In the orientation tuning results discussed above (figure 7) we have fixed the sharpness parameter *σ* of the stimulation protocol at an arbitrary value of 0.5. This value have generated orientation tuning surprisingly well matched against the tuning due to visual stimulation as is. However, prosthetic visual application might require fine-tuning or changing the orientation tuning width, for example due to differences across species or retinotopic eccentricity. In figure 8 we have explored range of values of the sharpness parameter, showing that systematic increase or decrease of the the parameter value leads to corresponding sharpening or broadening of the orientation tuning of the induced cortical responses (figure 8BD). We can also see that we have been lucky with our arbitrary choice of the parameter value in results presented in figure 7, having chosen value close to the optimum for our model based on cat/macaque physiology. Interestingly, despite the very broad range of values of the *σ* parameter explored in figure 8, the range of resulting orientation tuning width of cortical responses is rather narrow. This again points to our proposed hypothesis that the recurrent intra-cortical processing tends to nudge the resulting activation patterns towards those due to similar visual stimulation. We have thus demonstrated that the orientation tuning induced by proposed stimulation protocol has room for fine-tuning for specific circumstances, which in the case of clinical application, could be part of perceptually driven calibration process for given subject.

**Figure 8:**
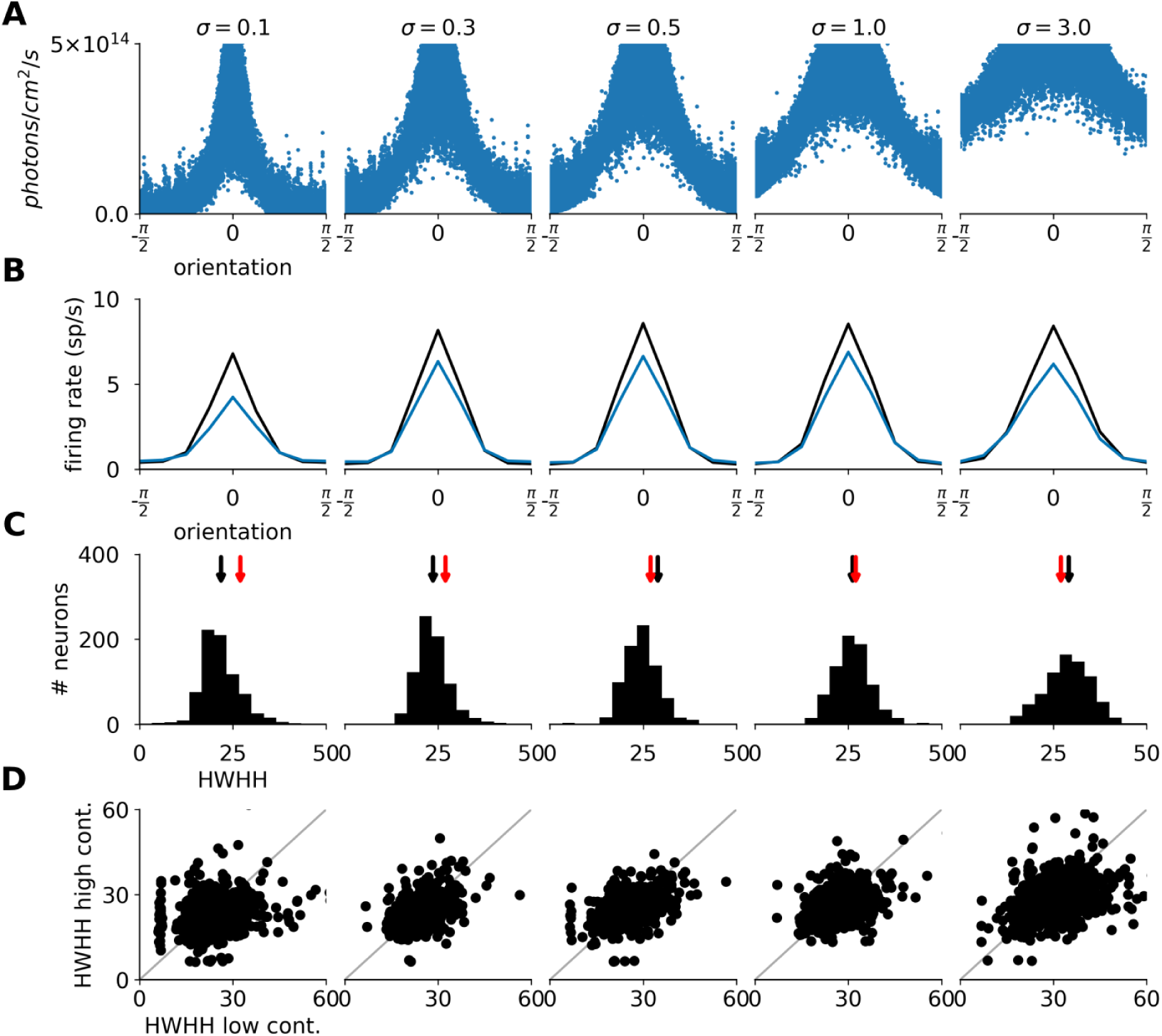
Width of orientation tuning as a function of the sharpness parameter of the optogenetic stimulation protocol. Each column corresponds to cortical optogenetic stimulation simulation where different parameters of the stimulation orientation sharpness *σ* was used. (A) The illumination intensity at the cell body as a function of the orientation preference of the given neuron. (B) The orientation tuning curves centered and averaged across all recorded neurons. (C) The scatter plot showing the orientation tuning width measured as HWHH at minimum and maximum contrast. (D) The histogram of tuning width measured as HWHH at maximum contrast. The black arrows on top mark the mean of the distribution. The red arrows mark the mean HWHH of the natural vision condition.

## 5 Statistical properties of the opogenetically evoked cortical activity

In the previous section we have shown that very good match between the response rates of visually and corresponding optogenetically evoked activity can be achieved. It can however be expected that properties of neural signals beyond that of mean response rate can have implications for perceptual outcomes. It is thus important to assess neural signals under optogenetic activation in greater detail. In the following we will focus on comparing basic statistical properties between the natural vision and optogenetic conditions both at the level of spikes but also membrane potential and excitatory and inhibitory conductances. For analysis in the section only neurons with orientation preference matching or orthogonal to the presented stimulus were analyzed.

A level of synchronization among nearby neurons during evoked activity has been demonstrated in primary visual cortex [75]. In figure 9A we show the synchrony among excitatory model neurons in layer 2/3 measured as the mean Pearson correlation of PSTH (binned at 10ms). As we can see the natural vision condition shows higher synchronization among neurons than the optogenetic stimulations. This is expected, as processing in thalamus and layer 4, up-stream from layer 2/3 and not engaged during light stimulation condition, can already induce some level of synchronization which can be further amplified in layer 2/3. Next figure 9A shows the ratio of excitation and inhibition during grating stimulation.

**Figure 9:**
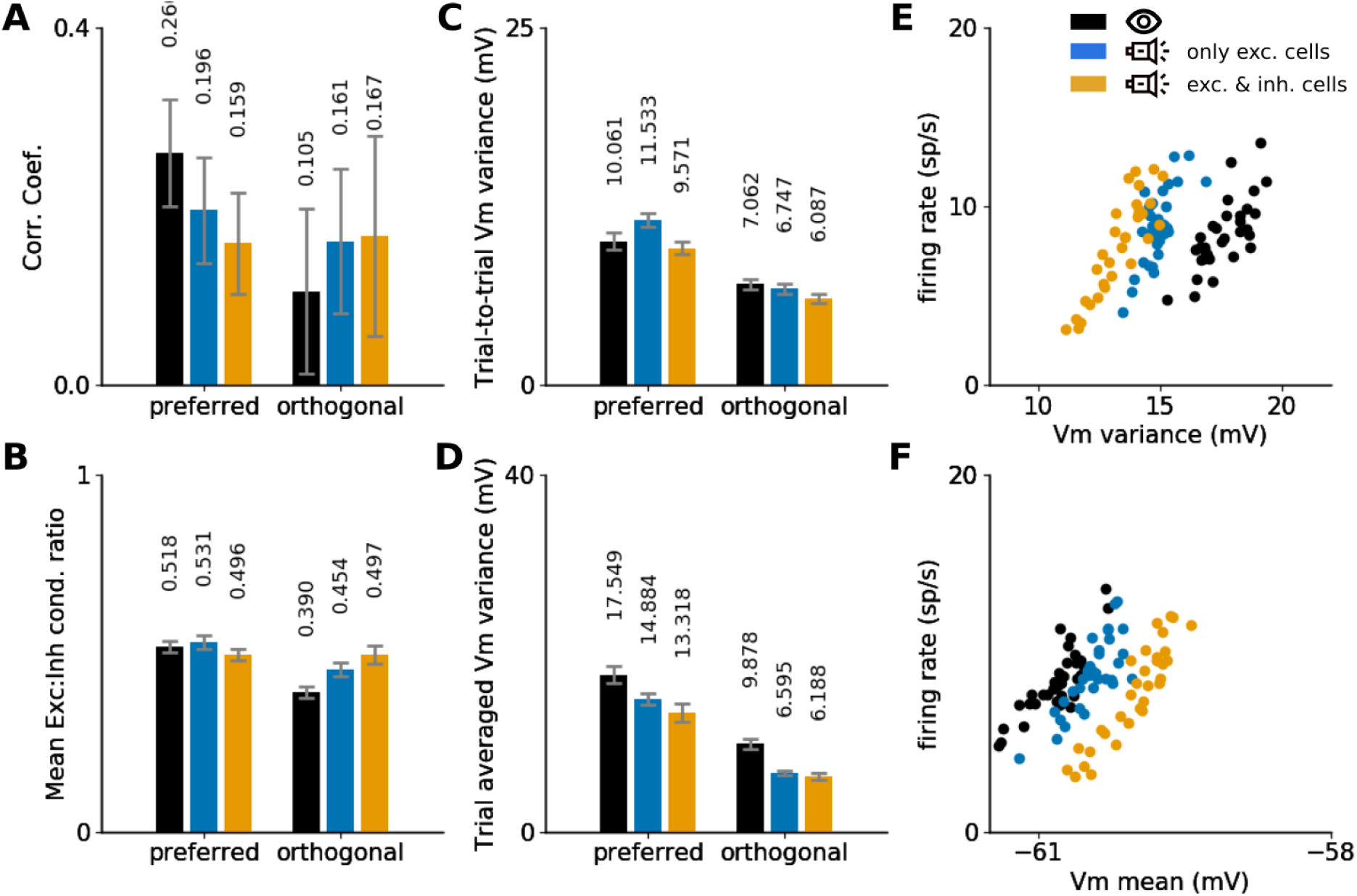
Visually and prosthetically evoked response statistics. (A-D) Response statistics at preferred (left bar group) and orthogonal (right bar group) orientation of the grating in the three examined conditions: normal vision (black), light stimulation with excitatory cells transfected (blue) or both exc. and inh. cells transfected (orange). (A) The mean Pearson correlation between the PSTH (binned at 10ms) of all pairs of recorded excitatory cells. (B) The ratio of mean excitatory and inhibitory conductance during presentation of grating averaged across all recorded cells. (C) The mean trial-to-trial variability of membrane potential during stimulation across the recorded cells. (D) Variability of Vm, averaged over the duration of the stimulus presentation and across over all recorded cells. (E-F) The relationship between variance (E) and mean (F) of the membrane potential and the response rate of recorded excitatory cells to the stimulation with grating of preferred orientation.

Interestingly, during the preferred orientation, neurons across all conditions show very similar ratios (*±* 5%). This is somewhat surprising, as for example the condition where only excitatory neurons are light-activated, one could expect bias towards excitation in the network. This again emphasizes that the recurrent cortical circuitry converges to similar operating regime during broad range of input scenarios. For the orthogonal orientation the one noticeable difference is that the natural vision condition has lower exc. to inh. ratio than the optogenetic conditions, indicating that visual stimulation engages more effectively lateral inhibition then light stimulation.

Next we looked at trial-to-trial (figure 9C) and trial averaged (figure 9D) variance of the Vm. First we can notice significant decrease of both quantities at orthogonal orientation (figure 9) as would be expected due to lower response rate. The most noticeable difference between the natural vision and light stimulation conditions is that the trial-averaged variance of the membrane potential is significantly higher in the natural vision condition, both at preferred and orthogonal stimulus configurations. This indicates that the visual stimuli induce more repeatable trial-to-trial responses than the light stimulation.

Ultimately it is the spike response of the cells that will determine their impact on perception. There are however two basic principles by which changes in membrane potential can affect the rate of spike generation: by changes in the mean (the mean driven regime) or by changes in the magnitude of variability that changes the likelyhood of crossing the spike generation threshold (fluctuation driven regime) [71, 47]. To compare the operating regime between the natural vision and optogenetic conditions we have plotted the spike rate response as a function of the mean (figure 9E) and the variance (figure 9G) of the membrane potential. As expected in all conditions both the mean and variance of membrane potential are positively correlated with spike rate, however major differences between the absolute levels of these measures are present between the different conditions. The natural vision condition remains less depolarized during the evoked activity than the optogenetic conditions, but instead relies on higher variability of the membrane potential to generate the same levels of response rates. Interestingly, the optogenetic condition with only excitatory light-sensitive neurons is closer to the natural vision condition, and notice that in fact in all the measures examined in this section the difference between the natural vision and excitatory only activation is smaller or the same compared to the condition where both cell types are activated. Overall these results show, that even though at the level of mean spike rate responses very good match between natural vision and optogenetic stimulation can be achieved (see section 4), important differences can remain at the sub-threshold level. It will therefore be important to quantify these differences in in-vivo, and determine in future experiments how much can such differences impact the perceptual outcomes. Our simulations also suggest, that targeting of only excitatory cells is preferable for cortical optogenetic prosthesis.

## 6 Impact of the density of the MLEE on the induction of orientation tuning

In all the simulations discussed so far we have assumed an MLEE with pitch and diameter of the individual light emitting elements of 10 *µm*. This value is rather at the extreme of what current technology can deliver and we used it to assess the full potential of optogentic prosthesis with current technology. However, as we have demonstrated in previous sections, our stimulation strategy rests upon the presence of topologically organized functional representations of the visual stimulus along cortical surface. It should be emphasized that this is not only our choice, but a necessity due to the geometrical constrains of the light illumination from MLEE, whereby dispersion of light in the neural tissue leads to loss of resolution of the light pattern with increasing cortical depth. On the other hand the smooth representation of visual feature along cortical surface implies that MLEE with greater pitch and size of individual light emitting elements could still be effective at functionally specific activation of primary visual cortex.

We thus set out to explore this possibility by running a series of optogenetic simulations with increasingly large diameter (and consequently pitch) of the light emitting elements. As we can see in figure 10B, the optogenetic stimulation can accurately match the orientation tuning due to visual stimulation up to the size of individual light emitting elements of 100 *µm*, beyond which the optogenetic response similarity starts to deteriorate. The contrast-invariance of the tuning is also largely maintained (figure 10C). We can again see that despite the fact that the orientation specific sharpness of the illumination degrades with increasing element diameter (figure 10A) the intra-cortical connectivity is able to sharpen the illumination pattern greatly to produce orientation tuning close to that due to natural vision over surprisingly broad range of element sizes (figure 10B,D). This finding indicates that utilization of lower resolution MLEEs can offer sufficient resolution of light patterns for vision restoration. This has important practical consequences, because lower resolution MLEEs are easier and consequently cheaper to manufacture, and crucially have greater potential to achieve sufficient light density, which is a major hurdle in manufacturing of MLEEs suitable for optogenetic stimulation.

**Figure 10:**
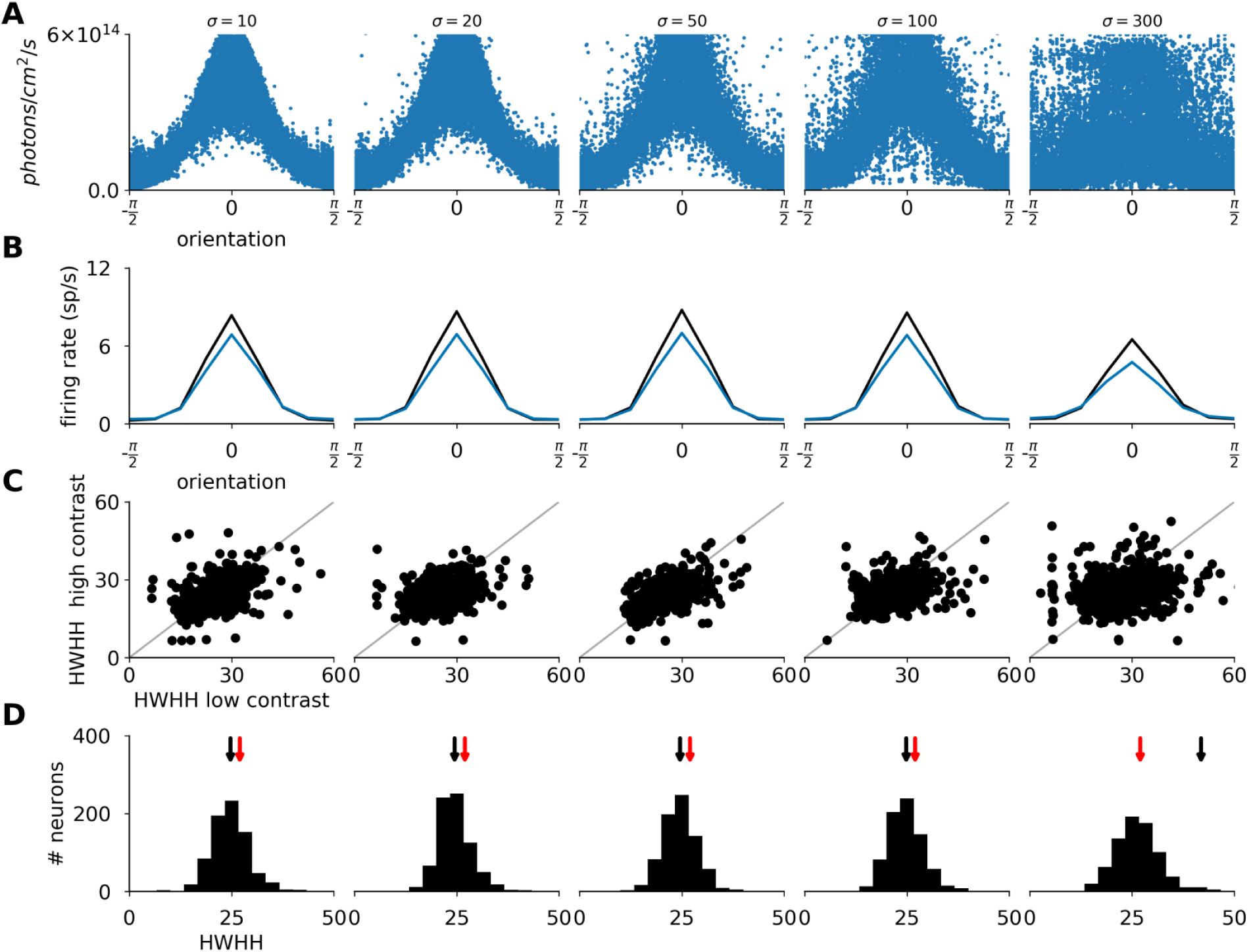
The impact of the diameter of individual light emitting elements on the induction of orientation tuning in V1. Each column corresponds to cortical optogenetic stimulation simulation with different size and pitch of light emitting elements. (A) The illumination intensity at the cell body as a function of the orientation preference of the given neuron. (B) The orientation tuning curves centered and averaged across all recorded neurons. (C) The scatter plot showing the orientation tuning width measured as HWHH at minimum and maximum contrast. (D) The histogram of tuning width measured as HWHH at maximum contrast. The black arrows on top mark the mean of the distribution. The red arrows mark the mean HWHH of the natural vision condition.

It is import to point out that in this study we focus only on a single of several features represented along cortical surface of primary visual cortex: the orientation of the stimulus. It is reasonable to expect that, as we will outline in greater detail in discussion, other visual features also topologically represented along cortical surface will be incorporated into the stimulation protocols in future. Some of these, such as retinotopy or occular dominance are represented at lower spatial frequency (in cortical surface coordinates) than orientation [74] and thus the above findings will hold. However, others, such as direction or disparity preference, are represented at higher spatial frequency and thus the minimum resolution of the MLEE would have to be reassessed for protocols targeting these visual features. To that end, the simulation environment presented in this study represents an efficient and flexible way to design and assess such series of increasingly complex stimulation protocols prior their in-vivo implementation.

## 7 Dicsussions

Cortical optogenetics based prosthetics are an emerging approach for vision restoration, that promises to mitigate several issues with previous technologies utilizing various forms of direct electrical stimulation. Undoubtedly, major technological development still needs to be undertaken to achieve the readiness of such prosthetic system for implantation in blind patients. Imagine, however, that the technology is ready. How would we go about stimulating the brain in such manner that it elicits useful perception of the visual world? Despite clearly being a central question of prosthetic vision, we are not aware of any study that attempted to directly answer this question. Because the relevant in-vivo experimental setups are not yet fully available, but some basic answers to this broad question will be needed to guide the in-vivo experiments that are expected in near future, we have set out to address this question via comprehensive computational modeling means.

Of course we do not intend to answer this question in its entirety. Concerted development in the field over next decades will be required for that, but this study does address some important initial concerns. We predict the intensity of illumination required for invocation of realistic levels of activity, and how will the required intensity change depending on which neural types are targeted by ChR transfection. We also provide guidance on the design of the stimulation hardware, for example showing how the size of the individual light emitting elements will effect the ability to evoke desired cortical active patterns. Our insights into how does the intra-cortical recurrent circuitry interact with the light-mediated input, how does the natural and prophetically evoked responses differ at sub-threshold level, how to translate between intensity of the stimulation and the contrast of the to be evoked stimulus will inform future more elaborate stimulations strategies. Last but not least, we provide here a ready to use stimulation protocol for induction of canonical visual stimuli, which is ideal for testing in early prosthetic experiments. Crucially, this stimulation strategy is accompanied by detailed prediction of what response should be expected. Of course it is likely that not all these predictions will be fulfilled, but by interpreting the experimental outcomes in the context of the models and by resolving any discrepancies that arise first in the model and thus gaining the understanding why they happened, will be instrumental for guiding the further development. We contend, that only trough such tight continuous interplay between modeling and experiments we can sufficient understanding of the cortico-prosthetic system to eventually achieve successful vision restoration.

### 7.1 Interaction of cortical circuitry with optogenetic stimulation

Given knowledge of the cortical responses pattern *R* elicited by presentation of stimulus *S* under natural vision, the straightforward way to approach the question of how to externally stimulate cortex to elicit responses similar to *R*, is to attempt to directly inject the response pattern *R* via the stimulation device (or its best possible approximation). In other words, to attempt to clamp the activity of the targeted cortical volume to its desired output. This approach, however, is guaranteed to work only if neurons were isolated entities. Quite contrary, individual neurons receive inputs via large number of synapses. This means that any externally imposed injection of activity into the cortical population will be further remodeled by the recurrent dynamics of the cortical network and the resulting activity pattern will presumably differ from that due to stimulation alone. On the other hand, one could follow the alternative hypothesis, that the optimal stimulation protocol should inject what would be the input to the target cortical population under normal vision, expecting that the cortical circuitry will reshape this simulated input the same way as the input due to normal vision and thus converge onto the same steady state response. Ultimately, it is not clear how significant factor will reshaping of injected activity via the intra-cortical circuitry be, especially in the context of various physical constraints of the prosthetic device which greatly limit the domain of spatio-temporal patterns illumination that can be achieved.

To advance our understanding of these issues, in this study we have compared the responses in V1 model between fully connected or disconnected conditions. We have found that taking the recurrent cortical interactions in consideration is crucial for arriving with correct predictions on how will cortex react to optogenetic stimulation, and consequently how to design an optimal stimulation protocol. Apart from differences of basic parameters such as excitability (section 3.2), which however can have major implications on the minimum specifications required of the stimulation device, we found that lateral cortical interactions sharpen the resulting patterns of activity in comparison to the light-induced pattern of depolarization, and consequently mitigate some of the systematic biases in the illumination patterns that arise due to the physical limitations of the stimulation technology. We observe analogous effects also when investigating the effect of sharpness of stimulation, observing much narrower range of tuning width of the resulting cortical responses when compared to the explored range of illumination tuning widths 4, and also when investigating the influence of light-emitting elements size (section 6). These findings are consistent with the idea, that it might be more appropriate to design prosthetic stimulation protocols such, that they mimic the functional input to the targeted cortical volume, rather than its output. In light of these observation we would like to propose following hypothesis: the intra-cortical recurrent circuitry tends to push externally induced cortical activation patterns towards similar proximate ecological (i.e. those that can arise due to plausible ecological visual stimulus) cortical response patterns. Overall these are encouraging findings, indicating that the intra-cortical circuitry can actually simplify the implementation of prosthetic vision, as they can mitigate some of the imprecisions in the illumination patterns, whether they are due to limitation to the stimulation strategy itself, or the various bio-physical constraints of the cortico-prosthetic system.

### 7.2 Future work

Being the first foray into modeling of cortical optogenetic stimulation, the present work has several limits that will need to be addressed in future. First, by focusing on the question of cortical network effects, we employ the much more computationally efficient point-neuron integrate and fire scheme as the model of neurons. These approaches have been shown to be effective at reproducing cortical processing [62, 11, 10] but important considerations might arise from the interaction between light illumination and the ChR channel distribution throughout the full neural morphology. Currently, computational resources are not available to perform simulations of the scale presented here that would take into account the full morphological detail of neurons. However, hybrid-schemes could be considered in the future, where for example the center of the model would be occupied by detailed morphological set of columns, but embedded within larger point-neuron network.

Another limitations of this study is that for now we have explored only the canonical stimulus class of drifting grating stimuli. This was a logical decision given that this is a first study of its kind, and this stimulus class is by far the best understood in terms of its V1 coding under normal vision condition, and thus easier to interpret under the new conditions we explore them here. As we can see range of useful insights that will inform future development of more elaborate and general stimulation protocols has been derived from the present work. But, of course, eventually more general stimulation protocols need to be developed if we are to achieve vision restoration. To this end in section 7.3 we propose a brief outline how the present work can be extended into full general purpose stimulation protocol. However, we would still like to note, that it still might be beneficial to study cortical stimulation in the context of more artificial stimuli, such as various protocol for studying contextual integration in V1, before moving to full ecological stimuli.

Another aspect of cortical processing that has not been explored here in greater details is the temporal dynamics of evoked responses. This is partially because representation of grating stimuli in complex cells largely reduce to a simple step depolarization function. However, as we show in figure 5, even for this temporally simple stimulus the cortical response in the natural vision condition show the characteristic onset and offset response transients, which our protocol did not take into account. Ultimately more work will have to be done to develop stimulation strategies that can accurately reproduce the stimulus evoked temporal dynamics of intact vision, especially in the context of more ecological visual stimuli. To this end, the advanced stimulation protocol, that we will describe in section 7.3, already offers a straightforward way to incorporate the canonical temporal response characteristics, such as the onset and offset dynamics.

### 7.3 Towards induction of arbitrary visual stimuli

In the present work we have only focused on a single canonical stimulus type in visual neuroscience: the sinusoidal gratings. This is because the responses of V1 to these stimuli are extremely well characterized, they are optimized to engage the most prominent functional property of V1 - orientation tuning, and their mathematical form with few parameters makes them natural starting point for investigation of stimulus-dependent phenomena. However, vision restoration obviously requires system that can induce perception of far more complex stimuli. We would like to point out that the work presented here is an essential step towards such vision restoration end-goal, goes beyond past stimulation protocols that were purely retinotopy based [61, 33, 66], and in fact forms basis for a straightforward extension into a protocol applicable to any visual stimulus. The present work enlightens the interaction between the light-induced input and recurrent network interactions, shows how to map stimulus contrast onto light stimulation intensity, shows that the cortical activations are responsive to changes in stimulation parameters in a predictable manner that allows for devising methods for fine-tuning the prospective stimulation protocols, and brings insights on the dependence of the opto-induced cortical activity patterns on the resolution of the light-stimulation array. All these findings should be highly informative for any future development of more complex cortical opto-stimulation protocols.

Let us now outline a straightforward path from the present work towards a stimulation protocol that can be applied to arbitrary visual stimuli. By only considering full-field stimuli of uniform orientation, here we have only considered the orientation domain of the cortical visual representation. If we would want to expand this consideration also into the retinotopic domain, a natural construct to consider is the ubiquitous gabor-shaped receptive field model [28] - i.e. the canonical model of V1 simple cells that binds the position and orientation preference of V1 neurons. Because we want to induce stimulus representation in layer 2/3 predominantly occupied by complex cells that are insensitive to the phase of the stimulus, we need to take one step further and consider an extension of the Gabor RF models for complex cells - the so called energy model[28]. The energy model is typically considered as a representation of single neurons, but because within a single orientation column the position and orientation preference of neurons varies little while in layer 2/3 the neurons are insensitive to phase, it is reasonable to assume that the energy models is in fact also a good approximation of the stimulus dependent response of the whole column. Following these consideration we propose following protocols:

1. Associate a energy model *µ*_*E*_ with each element *E* of the LED array, whose orientation and position is set to the pre-determined retinotopic and orientation preference of the cortical column below the element *E* and the remaining parameters are set to the know mean values at the given retinotopic eccentricity.
2. The activation of the LED element *E* is then proportional to the dot-product between *µ*_*E*_ and the visual stimulus to be induced.
3. To determine an appropriate mapping between the output of the RF model and the activation level of the LED element one can use analogous procedure as presented here in section 3.3.

This stimulation protocol is applicable to arbitrary stimulus, and can simply be extended to elementary temporal properties of V1 neural response by for example assuming space-time separability and expanding the RF model by a temporal filter reflecting the onset or offset dynamics. An interesting potential approach for setting the parameters of these filters would be to use reverse correlation approaches for their determination [68, 9] directly from data.

Furthermore, here we have restricted our consideration only to two stimulus feature selectivities present in V1 - the orientation and position. However, more such selectivities that are topologically represented along the cortical surface, and thus potentially exploitable by the optogenetic prosthetic setup considered here, are present in V1, including ocular dominance, ON-OFF dominance, direction, disparity, color and frequency preference. In principle, it would be straightforward to expand the proposed protocol to all these domains, providing they can be measured prior the implantation, as simple extensions of the RF model exist for all these features [21, 76, 2, 45]. It should be noted, however, that as more dimensions of the feature space are added the size of the cortical domains with similar properties across all these dimensions will shrink, and thus it would have to be determined if the opto-genetic prosthesis would have sufficient resolution to selectively engaged such higher-dimensional feature space of the cortical representation. Finally, the physical constraints of the considered opoto-genetic setup allowed us to consider only engagement of cortical representations that are topologically mapped onto cortical surface, however, the present work is informative and could be further expanded also to potential future technologies that facilitate activation of cortex at single-cell resolution (e.g. 2-or 3-photon imaging).

### 7.4 Consideration for application in blind subjects

The stimulation protocol presented in this study, as well as all the proposals for future extensions discussed in the previous section require the knowledge of the mapping of the neural stimulus feature preferences onto the cortical surface. During testing of the visual prosthesis in animal models, this information can be acquired by employing well established imaging techniques, such as intrinsic optical imaging [17], to measure these stimulus feature preferences directly using visual stimuli in an intact-vision condition. In this setting the ‘blind’ condition is induced later, and possibly reversibly, via surgical or chemical intervention. However, what about the ultimate goal of application to blind patients, where the intact-vision condition and thus visual stimulus dependent mapping is not available? The previous cortical visual prosthetic attempts solved this problem via a crude sequential calibration, based on reported perceptual outcomes in response to activation of individual or local groups of stimulation elements [33, 54]. Since past visual prosthetic devices only targeted single dimension of the visual feature representation - the retinotopic position, this procedure was relatively straightforward. The individual stimulation elements of the prosthesis were activated one by one, and for each activation the patient had to report the position of the elicited phosphene (or lack of it) in his visual field. Simple aids have been built to help sufficiently accurate reporting of the posphene position in the subjects visual field [54]. Furthermore, past empirical evidence have shown that this calibration procedure has to be repeated regularly, due to bio-physical processes [33, 54], that degrade the validity of the mapping over time.

However, this poses a problem for the optogenetic prosthetic approach proposed here, which would render such simple sequential calibration either highly impractical or outright impossible, due to the huge number of stimulation elements in the high-resolution LED array and the combinatorial explosion due to higher dimensional feature space. We would like to propose here that because thanks to the high-resolution opto-genetic prosthesis one has much finer control over the cortical activation, and because of the topological organization of the stimulus features targeted in this proposal, one could devise much more efficient calibration protocol. Assuming a rapid approximate identification of the retinotopic area that the LED array implant spans in the subject, that could for example be done using similar simple approach as in the past, one could identify the expected spatial frequency (along cortical surface) of the topological mapping of the given feature on the cortical surface at the pre-determined retinotopic eccentricity using previously established human functional atlas. One can than exploit the regularity of the functional topological mapping of the individual visual features (e.g. orientation) to radically reduce the number of stimulus presentations and associated reporting of perceptual outcomes required to determine the mapping of the given feature onto the cortical surface under the implant. Furthermore, one can repeat this process one feature at a time, thus braking the combinatorial augmentation. Of course, number of complications can be expected, for example distortions in the uniformity of functional representations due to presence of anatomical features such as vasculature, and ultimately only empirical experiments will resolve this issue, but we believe the proposed direction of investigation is worthwhile.

### 7.5 Broader impacts

The models and computational tools presented here allow us to examine the relationship between specific configurations of light delivered to cortex and the resulting spatio-temporal pattern of activity evoked in the simulated cortical circuitry. We have used them to advance our understanding of cortical stimulation under physiological conditions and to design a protocol for translation of specific class of visual stimuli to LED array activation pattern. In future, this same approach can be applied to further development of stimulation protocols, to screen numerous aspects of potential stimulation strategies in advance before the entire technological prosthetic stack is fully developed, and selecting only the most promising stimulation protocols for future testing in in-vivo animal models. This way one can inform the specification of future hardware components, as wells as informing the type of experiments that will be most informative in in-vivo testing, thus accelerating the development and reducing the usage of animal experimentation. In similar fashion, these computational tools could find use throughout the field of cortical based prosthetics, and could be adapted to address other cortical areas, sensory modalities and other stimulation techniques.

Finally, the high-fidelity modeling methodology developed in this study, together with the assumed prosthetic system, have potential to significantly advance our understanding of processing in the early visual cortex and beyond. On one hand, by directly probing neural activity, neuroscience has been effective in gaining insights into how visual information is coded in early sensory corticies and how is this coding implemented in neural substrate [57, 58]. On the other hand, using psychophysics, substantial insights into the principles of perception have been obtained [63]. However, bridging the gap between such low- and high level representations - i.e. understanding how low level coding leads to perception, and linking perceptual phenomena to neural substrate - remains elusive. In recent years optogenetic based brain stimulation in behaving rodents have attempted to addressee these issues [63], however the limited ability to induce complex activity patterns and poor behavioral potential of rodents prevented major breakthroughs.

The combination of high-fidelity modeling approach presented here with the optogenetic prosthesis could lead to paradigm shift in how neural basis of perception can be studied. Stimulation protocols reflecting specific properties of cortical sensory coding, and deep understanding how they interact with the cortical circuitry can be developed in the presented computational tools. Specific physiological and crucially perceptual predictions can be formulated, and subsequently tested in primate optogenetic model, utilizing behavioral psychohpysical experimental paradigms. This way a an unprecedented depth of understanding of the cortical processing that bridges the sensory coding, it’s implementation to neural circuitry and its perceptual outcome could be obtained.

## Supporting Information

**Figure S1:**
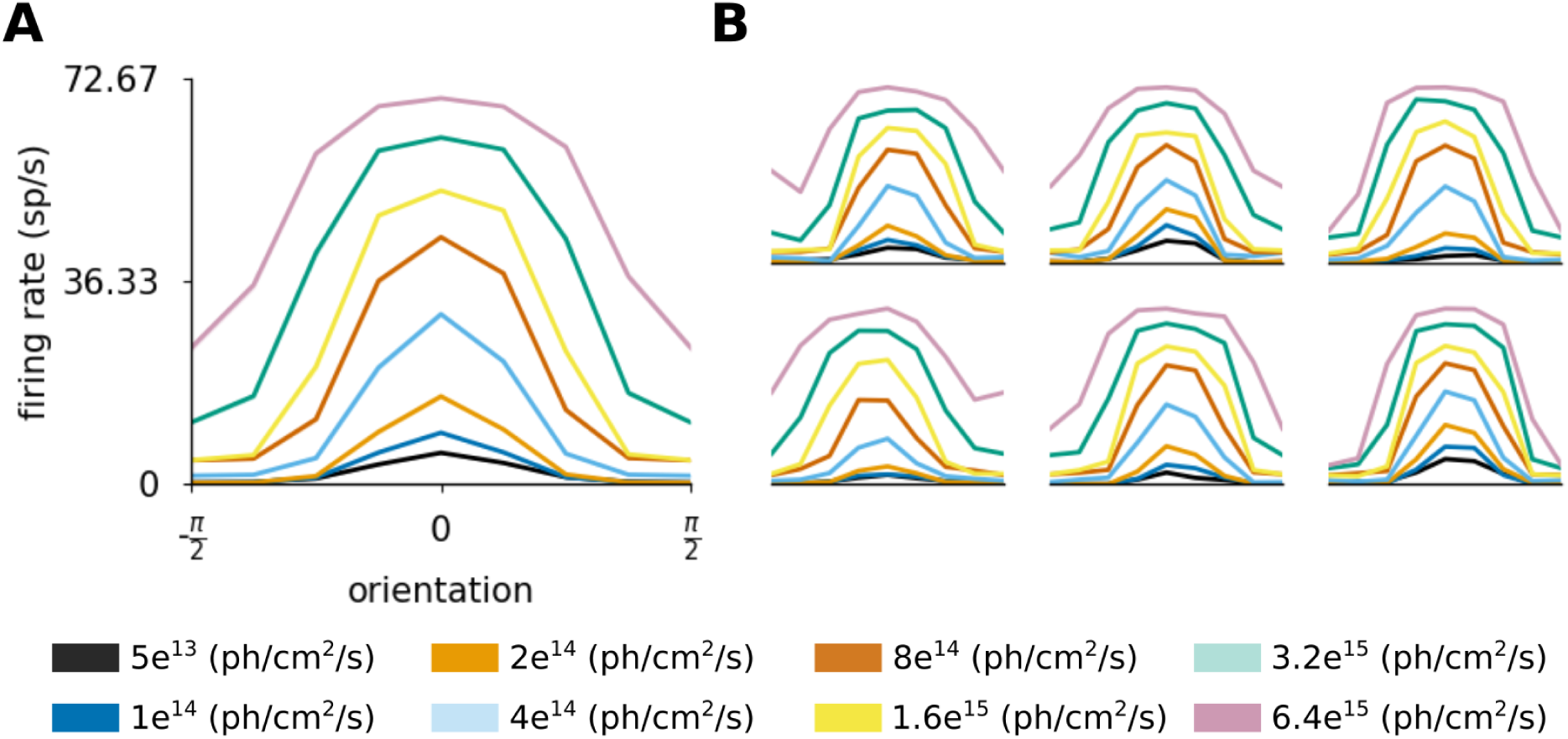
Orientation tuning of model neurons in the cortical stimulation condition, when only excitatory neurons are light sensitive. Orientation tuning curves are shown at different intensities of overall light stimulation (see legend). (A) mean of centered orientation tuning curves across all recorded neurons. (B) single cell tuning curves of 6 randomly selected neurons. The legend refers to maximum photon flux at the surface of the virtual implant.

## References

[1] L. F. Abbott. Synaptic Depression and Cortical Gain Control. Science, 275(5297):221–224, 1997.

[2] E. H. Adelson and J. R. Bergen. Spatiotemporal energy models for the perception of motion. Journal of the Optical Society of America A, 2(2):284, feb 1985.

[3] H. J. Alitto. Influence of Contrast on Orientation and Temporal Frequency Tuning in Ferret Primary Visual Cortex. Journal of Neurophysiology, 91(6):2797–2808, jun 2004.

[4] E. Allen and R. Freeman. Dynamic spatial processing originates in early visual pathways. The Journal of neuroscience, 26(45):11763–11774, 2006.

[5] A. Angelucci, J. B. Levitt, E. J. S. Walton, J.-M. Hupe, J. Bullier, and J. S. Lund. Circuits for local and global signal integration in primary visual cortex. The Journal of neuroscience : the official journal of the Society for Neuroscience, 22(19):8633–8646, oct 2002.

[6] Antolíkand J. J. A. Bednar. Development of Maps of Simple and Complex Cells in the Primary Visual Cortex. Frontiers in Computational Neuroscience, 5(April):1–19, 2011.

[7] J. Antolíkand A. A. Davison. Arkheia: Data Management and Communication for Open Computational Neuroscience. Frontiers in Neuroinformatics, 12:6, mar 2018.

[8] J. Antolíkand A. P. Davison. Integrated workflows for spiking neuronal network simulations. Frontiers in Neuroinformatics, 7(December):1–15, 2013.

[9] J. Antolík, S. S. B. Hofer, J. A. J. Bednar, and T. T. D. Mrsic-Flogel. Model Constrained by Visual Hierarchy Improves Prediction of Neural Responses to Natural Scenes. PLoS Computational Biology, 12(6):1–22, 2016.

[10] J. Antolík, C. Monier, A. Davison, and Y. Frégnac. A comprehensive data-driven model of cat primary visual cortex. bioRxiv, page 416156, jan 2019.

[11] A. Arkhipov, N. W. Gouwens, Y. N. Billeh, S. Gratiy, R. Iyer, Z. Wei, Z. Xu, R. Abbasi-Asl, J. Berg, M. Buice, N. Cain, N. da Costa, S. de Vries, D. Denman, S. Durand, D. Feng, T. Jarsky, J. Lecoq, B. Lee, L. Li, S. Mihalas, G. K. Ocker, S. R. Olsen, R. C. Reid, G. Soler-Llavina, S. A. Sorensen, Q. Wang, J. Waters, M. Scanziani, and C. Koch. Visual physiology of the layer 4 cortical circuit in silico. PLOS Computational Biology, 14(11):e1006535, 2018.

[12] Y. Banitt, K. A. C. Martin, and I. Segev. A Biologically Realistic Model of Contrast Invariant Orientation Tuning by Thalamocortical Synaptic Depression. Journal of Neuroscience, 27(38):10230–10239, sep 2007.

[13] C. Beaulieu and M. Colonnier. A laminar analysis of the number of round asymmetrical and flat symmetrical synapses on spines, dendritic trunks, and cell bodies in area 17 of the cat. Journal of Comparative Neurology, 231(2):180–189, 1985.

[14] C. Beaulieu and M. Colonnier. Number of neurons in individual laminae of areas 3B, 4*γ*, and 6a*α* of the cat cerebral cortex: A comparison with major visual areas. Journal of Comparative Neurology, 279(2):228–234, 1989.

[15] C. Beaulieu, Z. Kisvarday, P. Somogyi, M. Cynader, and A. Cowey. Quantitative distribution of gaba-immunopositive and - immunonegative neurons and synapses in the monkey striate cortex (area 17). Cerebral Cortex, 2(4):295–309, 1992.

[16] T. Binzegger. A Quantitative Map of the Circuit of Cat Primary Visual Cortex. Journal of Neuroscience, 24(39):8441–8453, sep 2004.

[17] G. Blasdel. Orientation selectivity, preference, and continuity in monkey striate cortex. The Journal of Neuroscience, 12(8):3139–3161, 2018.

[18] V. Bonin. The Suppressive Field of Neurons in Lateral Geniculate Nucleus. Journal of Neuroscience, 25(47):10844–10856, nov 2005.

[19] V. Bringuier, F. Chavane, L. Glaeser, and Y. Frénac. Horizontal Propagation of Visual Activity in the Synaptic Integration Field of Area 17 Neurons. Science, 283(5402):695–699, 1999.

[20] J. M. Budd and Z. F. Kisvárday. Local lateral connectivity of inhibitory clutch cells in layer 4 of cat visual cortex (area 17). Experimental Brain Research, 140(2):245–250, sep 2001.

[21] P. Buzás, P. Kóbor, Z. Petykó, I. Telkes, P. R. Martin, and L. Lénárd. Receptive field properties of color opponent neurons in the cat lateral geniculate nucleus. The Journal of neuroscience : the official journal of the Society for Neuroscience, 33(4):1451–61, jan 2013.

[22] P. Buzás, K. Kovács, A. S. Ferecskó, J. M. Budd, U. T. Eysel, and Z. F. Kisvárday. Model-based analysis of excitatory lateral connections in the visual cortex. Journal of Comparative Neurology, 499(6):861–881, ec 2006.

[23] M. Carandini. Do We Know What the Early Visual System Does? Journal of Neuroscience, 25(46):10577–10597, 2005.

[24] J. A. Cardin, L. A. Palmer, and D. Contreras. Stimulus Feature Selectivity in Excitatory and Inhibitory Neurons in Primary Visual Cortex. Journal of Neuroscience, 27(39):10333–10344, 2007.

[25] B. Chapman, M. P. Stryker, and T. Bonhoeffer. Development of orientation preference maps in ferret primary visual cortex. The Journal of neuroscience : the official journal of the Society for Neuroscience, 16(20):6443–53, oct 1996.

[26] N. M. da Costa and K. A. C. Martin. How Thalamus Connects to Spiny Stellate Cells in the Cat’s Visual Cortex. Journal of Neuroscience, 31(8):2925–2937, feb 2011.

[27] L. da Cruz, J. D. Dorn, M. S. Humayun, G. Dagnelie, J. Handa, P.-O. Barale, J.-A. Sahel, P. E. Stanga, F. Hafezi, A. B. Safran, J. Salzmann, A. Santos, D. Birch, R. Spencer, A. V. Cideciyan, E. de Juan, J. L. Duncan, D. Eliott, A. Fawzi, L. C. Olmos de Koo, A. C. Ho, G. Brown, J. Haller, C. Regillo, L. V. Del Priore, A. Arditi, and R. J. Greenberg. Five-Year Safety and Performance Results from the Argus II Retinal Prosthesis System Clinical Trial. Ophthalmology, 123(10):2248–2254, oct 2016.

[28] J. G. Daugman. Uncertainty relation for resolution in space, spatial frequency, and orientation optimized by two-dimensional visual cortical filters. Journal of the Optical Society of America. A, Optics and image science, 2(7):1160–9, jul 1985.

[29] G. DeAngelis, I. Ohzawa, and R. Freeman. Spatiotemporal organization of simple-cell receptive fields in the cat’s striate cortex. I. General characteristics and postnatal development. Journal of Neurophysiology, 69(4):1091, 1993.

[30] K. Deisseroth. Optogenetics : 10 years of microbial opsins in neuroscience. Nature Neuroscience, 18(9):1213–1225, 2015.

[31] A. Destexhe, Z. F. Mainen, and T. J. Sejnowski. Synthesis of models for excitable membranes, synaptic transmission and neuromodulation using a common kinetic formalism. Journal of computational neuroscience, 1(3):195–230, 1994.

[32] W. H. Dobelle. Introduction to sensory prostheses for the blind and deaf. Transactions - American Society for Artificial Internal Organs, 20 B:761–4, 1974.

[33] W. H. Dobelle. Artificial Vision for the Blind by Connecting a Television Camera to the Visual Cortex State of the Art. ASAIO Journal, 2000.

[34] I. M. Finn, N. J. Priebe, and D. Ferster. The emergence of contrast-invariant orientation tuning in simple cells of cat visual cortex. Neuron, 54(1):137–52, apr 2007.

[35] Y. Frénac. Reading out the synaptic echoes of low-level perception in V1. In A. Fusiello, V. Murino, and R. Cucchiara, editors, Lecture Notes in Computer Science (including subseries Lecture Notes in Artificial Intelligence and Lecture Notes in Bioinformatics), volume 7583 of Lecture Notes in Computer Science, pages 486–495. Springer, 2012.

[36] M.-O. Gewaltig and M. Diesmann. NEST (NEural Simulation Tool). Scholarpedia, 2(4):1430, 2007.

[37] C. D. Gilbert. Laminar differences in receptive field properties of cells in cat primary visual cortex. J Physiol, 268:391–421, 1977.

[38] J. A. Hirsch and L. M. Martinez. Laminar processing in the visual cortical column. Current Opinion in Neurobiology, 16(4):377–384, aug 2006.

[39] D. H. Hubel and T. N. Wiesel. Receptive fields, binocular interaction and functional architecture in the cat’s visual cortex. The Journal of Physiology, 160(1):106–154, jan 1962.

[40] D. H. Hubel and T. N. Wiesel. Sequence regularity and geometry of orientation columns in the monkey striate cortex. The Journal of comparative neurology, 158(3):267–93, ec 1974.

[41] S. L. Jacques. Optical properties of biological tissues: a review. Physics in Medicine and Biology, 58(11):R37–R61, jun 2013.

[42] E. L. Jocoy. Dissecting the contribution of individual receptor subunits to the enhancement of N-methyl-D-aspartate currents by dopamine D1 receptor activation in striatum. Frontiers in Systems Neuroscience, 5(February):1–26, 2011.

[43] J. P. Jones. The two-dimensional spatial structure of simple receptive fields in cat striate cortex. Journal of Neurophysiology, 58(6):1187–1211, ec 1987.

[44] H. Ko, S. B. Hofer, B. Pichler, K. A. Buchanan, P. J. Sjöström, and T. D. Mrsic-Flogel. Functional specificity of local synaptic connections in neocortical networks. Nature, 473(7345):87–91, apr 2011.

[45] J. Kremkow, J. Jin, S. J. Komban, Y. Wang, R. Lashgari, X. Li, M. Jansen, Q. Zaidi, and J.-M. J.-M. Alonso. Neuronal nonlinearity explains greater visual spatial resolution for darks than lights. Proceedings of the National Academy of Sciences of the United States of America, 111(8):3170–5, feb 2014.

[46] J. Kremkow, L. U. Perrinet, C. Monier, J.-M. Alonso, A. Aertsen, Y. Frénac, and G. S. Mason. Push-Pull Receptive Field Organization and Synaptic Depression: Mechanisms for Reliably Encoding Naturalistic Stimuli in V1. Frontiers in Neural Circuits, 10(May), 2016.

[47] A. Kuhn, A. Aertsen, and S. Rotter. Neuronal integration of synaptic input in the fluctuation-driven regime. The Journal of neuroscience : the official journal of the Society for Neuroscience, 24(10):2345–56, mar 2004.

[48] P. M. Lewis, H. M. Ackland, A. J. Lowery, and J. V. Rosenfeld. Restoration of vision in blind individuals using bionic devices A review with a focus on cortical visual prostheses. Brain Research, 1595:51–73, 2015.

[49] Z. Liang, W. Shen, C. Sun, and T. Shou. Comparative study on the offset responses of simple cells and complex cells in the primary visual cortex of the cat. Neuroscience, 156(2):365–373, 2008.

[50] H. Markram, M. Toledo-Rodriguez, Y. Wang, A. Gupta, G. Silberberg, and C. Wu. Interneurons of the neocortical inhibitory system. Nature Reviews Neuroscience, 5(10):793–807, oct 2004.

[51] H. Markram, Y. Wang, and M. Tsodyks. Differential signaling via the same axon of neocortical pyramidal neurons. Proc Natl Acad Sci U S A, 95(9):5323–8., apr 1998.

[52] J. H. Maunsell, G. M. Ghose, J. A. Assad, C. J. Mcadams, C. E. Boudreau, and B. D. Noerager. Visual response latencies of magnocellular and parvocellular LGN neurons in macaque monkeys. Visual Neuroscience, 16(1):1–14, 1999.

[53] C. Monier, J. Fournier, and Y. Frénac. In vitro and in vivo measures of evoked excitatory and inhibitory conductance dynamics in sensory cortices. Journal of Neuroscience Methods, 169(2):323–365, apr 2008.

[54] J. Naumann. Search for Paradise: A Patient’s Account of the Artificial Vision Experiment. Xlibris Corporation, 2012, 2012.

[55] L. G. Nowak, M. V. Sanchez-Vives, and D. A. McCormick. Lack of orientation and direction selectivity in a subgroup of fast-spiking inhibitory interneurons: Cellular and synaptic mechanisms and comparison with other electrophysiological cell types. Cerebral Cortex, 18(5):1058–1078, may 2008.

[56] O. Ohana, H. Portner, and K. A. C. Martin. Fast recruitment of recurrent inhibition in the cat visual cortex. PLoS ONE, 7(7), 2012.

[57] B. A. Olshausen. 20 Years of Learning About Vision: Questions Answered, Questions Unanswered, and Questions Not Yet Asked. In 20 Years of Computational Neuroscience, pages 243–270. Springer, 2013.

[58] B. A. Olshausen and D. J. Field. How Close Are We to Understanding V1? Neural Computation, 17(8):1665–1699, 2005.

[59] J. Papaioannou and A. White. Maintained activity of lateral geniculate nucleus neurons as a function of background luminance. Experimental Neurology, 34(3):558–566, 1972.

[60] X. Pei, T. R. Vidyasagar, M. Volgushev, and O. D. Creutzfeldt. Receptive field analysis and orietation selectivity of postsynaptic potentials of simple cells in cat visual cortex. The Journal of Neuroscience, 14(November):7130–7140, 1994.

[61] J. S. Pezaris, R. Clay Reid, and R. C. Reid. Demonstration of artificial visual percepts generated through thalamic microstimulation. PNAS, 104(18):7670–7675, 2007.

[62] A. V. Rangan, L. Tao, G. Kovacic, and D. Cai. Multiscale modeling of the primary visual cortex. IEEE Engineering in Medicine and Biology, 28(3):19–24, 2009.

[63] J. Read. The place of human psychophysics in modern neuroscience. Neuroscience, 296:116–129, jun 2015.

[64] D. L. Ringach, R. M. Shapley, and M. J. Hawken. Orientation selectivity in macaque V1: diversity and laminar dependence. The Journal of neuroscience : the official journal of the Society for Neuroscience, 22(13):5639–51, jul 2002.

[65] Q. Sabatier, C. Joffrois, G. Gauvais, J. Chavas, D. Pruneau, S. Picaud, and R. Benosman. Modeling the Electro-chemical Properties of Microbial Opsin ChrimsonR for Application to Optogenetics-based Vision Restoration. bioRxiv, page 417899, sep 2018.

[66] E. M. Schmidt, M. J. Bak, F. T. Hambrecht, C. V. Kufta, D. K. O’Rourke, and P. Vallabhanath. Feasibility of a visual prosthesis for the blind based on intracortical microstimulation of the visual cortex. Brain : a journal of neurology, 119 (Pt 2:507–22, apr 1996.

[67] R. K. Shepherd, M. N. Shivdasani, D. A. Nayagam, C. E. Williams, and P. J. Blamey. Visual prostheses for the blind. Trends in Biotechnology, 31(10):562–571, oct 2013.

[68] D. Smyth, B. D. B. Willmore, G. E. Baker, I. D. Thompson, and D. J. Tolhurst. The receptive-field organization of simple cells in primary visual cortex of ferrets under natural scene stimulation. The Journal of Neuroscience, 23(11):4746–59, jun 2003.

[69] A. Stepanyants, L. M. Martinez, A. S. Ferecsko, and Z. F. Kisvarday. The fractions of short- and long-range connections in the visual cortex. Proceedings of the National Academy of Sciences, 106(9):3555–3560, 2009.

[70] A. Stepanyants, L. M. Martinez, A. S. Ferecsko, and Z. F. Kisvarday. The fractions of short- and long-range connections in the visual cortex. Proceedings of the National Academy of Sciences, 106(9):3555–3560, jan 2009.

[71] L. Tao, D. Cai, D. W. McLaughlin, M. J. Shelley, and R. Shapley. Orientation selectivity in visual cortex by fluctuation-controlled criticality. Proceedings of the National Academy of Sciences of the United States of America, 103(34):12911–12916, aug 2006.

[72] T. W. Troyer, a. E. Krukowski, N. J. Priebe, and K. D. Miller. Contrast-invariant orientation tuning in cat visual cortex: thalamocortical input tuning and correlation-based intracortical connectivity. The Journal of Neuroscience, 18(15):5908–5927, aug 1998.

[73] R. J. Tusa, L. a. Palmer, and a. C. Rosenquist. The retinotopic organization of area 17 (striate cortex) in the cat. The Journal of comparative neurology, 177(2):213–35, jan 1978.

[74] M. Weliky, W. H. Bosking, and D. Fitzpatrick. A systematic map of direction preference in primary visual cortex. Nature, 379(6567):725–728, feb 1996.

[75] J. Yu and D. Ferster. Membrane Potential Synchrony in Primary Visual Cortex during Sensory Stimulation. Neuron, 68(6):1187–1201, ec 2010.

[76] Y. D. Zhu and N. Qian. Binocular receptive field models, disparity tuning, and characteristic disparity. Neural computation, 8(8):1611–41, nov 1996.

[77] A. Ziskind. Neurons in Cat Primary Visual Cortex cluster by degree of tuning but not by absolute spatial phase or temporal response phase. PhD Thesis, 2013.

